# An advanced sequence clustering and designation workflow reveals the enzootic maintenance of a dominant West Nile virus subclade in Germany

**DOI:** 10.1101/2022.10.05.509209

**Authors:** Pauline Dianne Santos, Anne Günther, Markus Keller, Timo Homeier-Bachmann, Martin H. Groschup, Martin Beer, Dirk Höper, Ute Ziegler

## Abstract

West Nile virus (WNV) is the most widespread arthropod-borne (arbo) virus and the primary cause of arboviral encephalitis globally. Members of WNV species genetically diverged and are classified into different hierarchical groups below species rank. However, the demarcation criteria for allocating WNV sequences into these groups remain individual, inconsistent, and the use of names for different levels of the hierarchical levels is unstructured. In order to have an objective and comprehensible grouping of WNV sequences, we developed an advanced grouping workflow using the “affinity propagation clustering”-algorithm and newly included the “agglomerative hierarchical clustering”-algorithm for the allocation of WNV sequences into different groups below species rank. In addition, we propose to use a fixed set of terms for the hierarchical naming of WNV below species level and a clear decimal numbering system to label the determined groups. For validation, we applied the refined workflow to WNV sequences that have been previously grouped into various lineages, clades, and clusters in other studies. Although our workflow regrouped some WNV sequences, overall, it generally corresponds with previous groupings. We employed our novel approach to the sequences from the WNV circulation in Germany 2020, primarily from WNV-infected birds and horses. Besides two newly defined minor (sub)clusters comprising only of three sequences each, subcluster 2.5.3.4.3c was the predominant WNV sequence group detected in Germany from 2018-20. This predominant subcluster was also associated with at least five human WNV-infections in 2019-20. In summary, our analyses imply that the genetic diversity of the WNV population in Germany is shaped by enzootic maintenance of the dominant WNV subcluster accompanied by sporadic incursions of other rare clusters and subclusters. Moreover, we show that our refined approach for sequence grouping yields meaningful results. Although we primarily aimed at a more detailed WNV classification, the presented workflow can also be applied to the objective genotyping of other virus species.

## 1. Introduction

Like other members of the genus *Flavivirus*, West Nile virus (WNV) has become a serious emerging zoonotic threat in Europe within the last decades (European Centre for Disease Prevention and Control n.d.; Kuno et al. 1998). The first known case of WNV-infection was reported in Uganda, Africa, in 1937 (Bardos et al. 1959; Smithburn et al. 1940). In the 1960s, the first occurrence of WNV in Europe was recognized due to neurological disorders in wild and domestic horses in France (Murgue et al. 2001). Around 30 years later, WNV caused the first severe outbreak of West Nile Fever (WNF) and West Nile Neuroinvasive Disease (WNND) in humans in Romania (Savage et al. 1999; Tsai et al. 1998). Since then, WNV has successfully established in various countries. Southern and eastern European countries were primarily affected by recurring WNV infections in humans, birds, and horses. The highest WNV activity in Europe was recorded in 2018 (Camp and Nowotny 2020; European Centre for Disease Prevention and Control 2019). Almost 90% of all locally acquired WNV human infections in Europe, with 166 fatal cases, were reported in Italy, Greece, and Romania (European Centre for Disease Prevention and Control 2019). In parallel to this large-scale epidemic in 2018, WNV-RNA positive birds and horses were confirmed for the first time in Germany (Ziegler et al. 2019). In 2019, a significant increase in WNV cases in birds and horses as well as the first five autochthonous WNV human infections in Germany were reported (Robert-Koch-Institut 2020; Ziegler et al. 2020). All prerequisites for endemic WNV circulation in Germany are fulfilled, including the proven vector competence of local mosquito populations (Holicki et al. 2020) and the detection of WNV genome-positive mosquito pools (Kampen et al. 2020; Ziegler et al. 2020).

WNV has a diverse host range and is widely distributed. Accordingly, members of this species are genetically diverse, allowing for the further subgrouping within the species. However, since the International Committee on Taxonomy of Viruses (ICTV) confines its responsibility to the designation and demarcation of viruses from realm to species ranks (ICTV 2020; Simmonds et al. 2017), neither a standard definition of criteria for subgrouping below the species rank nor defined designations for subgroups and their hierarchical arrangement exist. Therefore, designations for hierarchical ranks (*e.g.*, clade, cluster, sub-type, genotype) are often used inconsistently and interchangeably, leading to misunderstandings and uncertainties as more and more whole genomes of WNV are generated. Due to its aforementioned genetic diversity, up to nine lineages have been proposed for the species *West Nile virus* (Fall et al. 2017; Mencattelli et al. 2022; Pachler et al. 2014). The designation “lineage” is mostly based on monophyletic clustering of partial or whole genome WNV sequences in phylogenetic analyses (Fall et al. 2017; Perez-Ramirez et al. 2017). However, the lineage classification of WNV strains remains controversial (Perez-Ramirez et al. 2017). Further subgrouping within the lineages is conducted to organize viruses into a hierarchical system comprising of various arbitrarily defined and designated groups. Especially within and between members of WNV lineages 1 and 2 the designations are used inconsistently. Groups are usually defined based on branching into monophyletic groups from a common ancestor and members of groups may share common characteristics such as unique and fixed amino acid (aa) substitutions (Anez et al. 2013; Barzon et al. 2015; Chaintoutis et al. 2019; Davis et al. 2005; Di Giallonardo et al. 2016; Hadfield et al. 2019; May et al. 2011; McMullen et al. 2013; Ziegler et al. 2020). Monophyletic groups other than lineages are typically labelled using a letter, region of origin, or abbreviation of the region of origin (Fall et al. 2017; Kolodziejek et al. 2014; McMullen et al. 2013; Ravagnan et al. 2015; Zehender et al. 2017; Ziegler et al. 2019; Ziegler et al. 2020). Noteworthy, nomenclatures based on geographic origin may be misleading. For instance, a WNV sequence from Italy branched with Eastern European WNV lineage 2 sequences detected in Romania and Russia (Bakonyi and Haussig 2020; Ravagnan et al. 2015; Sikkema et al. 2020; Ziegler et al. 2020). Moreover, Ziegler and colleagues (Ziegler et al. 2020) mentioned in the study of the 2018-19 WNV epidemic in Germany that the label “Eastern German WNV Clade (EGC)”, designated to a group of WNV sequences from Germany, may not be a suitable designation because “the EGC can have developed in the wider southeastern and central European hemisphere and may have been translocated only later to Eastern Germany”. Hence, labels based on geographic origin may not suit the expanding geographic or undiscovered range of a WNV sequence group.

The described situation emphasizes the need for a systematic nomenclature and objective grouping of WNV sequences into hierarchical groups below the species rank. To subdivide WNV, we further developed the objective clustering workflow established by Fischer and colleagues (Fischer et al. 2018) who utilized the affinity propagation clustering (APC) algorithm (Frey and Dueck 2007) as implemented by Bodenhofer and colleagues (Bodenhofer et al. 2011). However, Fischer and colleagues found limitations of APC especially for the definition of the best suited number of clusters and therefore ultimately the definition of groups corresponding with phylogenetic analyses. To solve these issues, we refined the method to define a suitable number of groups while also incorporating agglomerative hierarchical clustering (AHC) (Bodenhofer et al. 2011) to address grouping of sequences into multiple hierarchical levels. In addition, we suggest a decimal numbering system for the hierarchical groups designated with the proposed unified and consistent labels within the WNV species. Finally, we provide an update on the WNV situation in birds and horses in Germany 2020 by applying the improved clustering workflow and our novel generic and consistent nomenclature.

## 2. Material and Methods

### 2.1 WNV screening of birds and horses

The nationwide wild bird surveillance program in Germany was established as an instantaneous reaction to the first Usutu virus (USUV) epizootic in 2011. This monitoring program became reputable also for the early detection of other zoonotic arboviruses, such as Sindbis virus and WNV. WNV infection in birds and horses is a notifiable animal disease in Germany ifdetected by RT-qPCR (real time quantitative polymerase chain reaction) and/or the identification of WNV-specific IgM in non-vaccinated horses by ELISA (enzyme-linked immunosorbent assay; i.e. detection of a recent WNV infection).

Samples from birds or horses (e.g. complete animals, organ samples, blood samples, and/or total RNA) were sent to the national reference laboratory for WNV at the Friedrich-Loeffler-Institut (FLI), Isle of Riems, Germany, by the regional veterinary laboratories of the German federal states, and by members of the nationwide wild bird surveillance program (for details about the members see (Ziegler et al. 2022)).

### 2.2 Ethical statement

Bird clinics, veterinarians, wild bird rescue centers and zoos provided bird carcasses for necropsy. In Germany, no specific permits are required to examine dead birds which have been submitted for necropsy. Horse clinics and veterinarians from the regional veterinary laboratories provided horse tissue samples collected in post-mortem examinations by pathological institutions. Residual blood material was available for one case originating from a WNV-infected bird, collected primarily for diagnostic purposes and for specific treatment and prognosis.

### 2.3 RNA extraction and RT-qPCR

Total RNA was extracted from tissue samples (brain, spleen, liver, spinal cord, and/or kidney) and frozen (-70 °C) coagulated blood samples (cruor). For the first RNA extraction, we applied the RNeasy Mini Kit (QIAGEN) according to the manufacturer’s instructions, followed by screenings for both WNV lineage 1 and 2 genomes using an RT-qPCR assay (Eiden et al. 2010).

### 2.4 Whole-genome sequencing

To cover areas with and without previous WNV cases, WNV RNA positive samples from 2020 (Table 1) were selected for whole-genome sequencing (WGS) primarily based on their geographical location and Cq values. In addition, samples from captive birds, wild birds, and horses from similar regions were included. These selected samples (Table 1) were subjected to a different RNA extraction protocol to ensure the acquisition of high-quality starting material for WGS. Briefly, each organ homogenate suspension (250 µl) was lysed in 750 µl TRIzol™ LS Reagent (Invitrogen) or approximately 30 mg tissue material homogenized in 1 ml TRIzol™ reagent via TissueLyser II (Qiagen) with a 5 mm steel bead for 2 min at 30 Hz. After phase separation, the aqueous phase was processed using the Agencourt® RNAdvance Tissue kit (Beckman Coulter) and the KingFisher Flex system (Thermo Fisher Scientific) according to the manufacturer’s instructions.

**Table 1.** Overview of WNV cases analyzed in this study. Sample numbers are used in Figures 4 and 7.

| Sample no. | Sample ID | Library ID | Sequence accession | Common English name | Scientific name | Collection date | Federal state<sup>1</sup> |
| --- | --- | --- | --- | --- | --- | --- | --- |
| 1 | 167/20 | lib04566 |  | Blue tit | *Cyanistes caeruleus* | 07.07.2020 | BE |
| 2 | 174/20 | lib04567 |  | Snowy owl | *Bubo scandiacus* | 12.07.2020 | TH |
| 3 | 192/20 | lib04568 |  | Snowy owl | *Bubo scandiacus* | 15.08.2020 | TH |
| 4 | 193/20 | lib04569 |  | Northern goshawk | *Accipiter gentilis* | 30.07.2020 | SN |
| 5 | 200/20 | lib04570 |  | Bohemian waxwing | *Bombycilla garrulus* | 11.08.2020 | BB |
| 6 | 203/20 | lib04571 |  | Little owl | *Athene noctua* | 18.08.2020 | BB |
| 7 | 206/20 | lib04572 |  | Northern goshawk | *Accipiter gentilis* | 08.08.2020 | BE |
| 8 | 214/20 | lib04573 |  | Unspecified flamingo | *Phoenicopterus sp.* | 19.08.2020 | TH |
| 9 | 218/20 | lib04574 |  | Unspecified flamingo | *Phoenicopterus sp.* | 18.08.2020 | ST |
| 10 | 339/20 | lib04575 |  | Chilean flamingo | *Phoenicopterus chilensis* | 25.08.2020 | BE |
| 11 | 252/20 | lib04576 |  | Eurasian jay | *Garrulus glandarius* | Aug. /Sept.2020 | TH |
| 12 | 283/20 | lib04577 |  | Snowy owl | *Bubo scandiacus* | 02.09.2020 | ST |
| 13 | 207/20 | lib04717 |  | Northern goshawk | *Accipiter gentilis* | 09.08.2020 | BE |
| 14 | 208/20 | lib04718 |  | Northern goshawk | *Accipiter gentilis* | 14.08.2020 | BB |
| 15 | 211/20 | lib04719 |  | Blue tit | *Cyanistes caeruleus* | August 2020 | BE |
| 16 | 194/20 | lib04720 |  | Blue tit | *Cyanistes caeruleus* | August 2020 | SN |
| 17 | 268/20 Nr. 1 | lib04721 |  | Little owl | *Athene noctua* | September 2020 | BB |
| 18 | 311/20 | lib04722 |  | Northern goshawk | *Accipiter gentilis* | Sept./Okt. 2020 | BE |
| 19 | 314/20 | lib04723 |  | Northern goshawk | *Accipiter gentilis* | Sept./Okt. 2020 | BE |
| 20 | 315/20 | lib04724 |  | Northern goshawk | *Accipiter gentilis* | Sept./Okt. 2020 | BE |
| 21 | 224/20 | lib04725 |  | Blue tit | *Cyanistes caeruleus* | August 2020 | BE |
| 22 | 282/20 | lib04726 |  | Snowy owl | *Bubo scandiacus* | 02.09.2020 | ST |
| 23 | 281/20 | lib04727 |  | Unspecified flamingo | *Phoenicopterus sp.* | September 2020 | ST |
| 24 | 340/20 | lib04728 |  | American flamingo | *Phoenicopterus ruber* | 22.08.2020 | BE |
| 25 | 196/20 | lib04729 |  | Unspecified buzzard | *Buteo sp.* | August 2020 | SN |
| 26 | 258/20 | lib04730 |  | Great tit | *Parus major* | Aug. /Sept.2020 | SN |
| 27 | 242/20 | lib04731 |  | Northern goshawk | *Accipiter gentilis* | August 2020 | BE |
| 28 | 245/20 | lib04732 |  | Unspecified flamingo | *Phoenicopterus sp.* | August 2020 | BE |
| 29 | 260/20 | lib04733 |  | Domestic canary | *Serinus canaria forma domestica* | September 2020 | ST |
| 30 | 261/20 | lib04734 |  | Chilean flamingo | *Phoenicopterus chilensis* | September 2020 | SN |
| 31 | 264/20 | lib04735 |  | Unspecified sparrow | *Passer sp.* | September 2020 | SN |
| 32 | 228/20 | lib04736 |  | Horse | *Equus caballus* | 02.09.2020 | SN |
| 33 | 173/20 | lib04737 |  | Alpine chough | *Pyrrhocorax graculus* | 12.07.2020 | ST |
| 34 | 216/20 | lib04738 |  | Blue tit | *Cyanistes caeruleus* | August 2020 | TH |
| 35 | 199/20 | lib04739 |  | Northern goshawk | *Accipiter gentilis* | 13.06.2020 | BE |
| 36 | 205/20 | lib04740 |  | Northern goshawk | *Accipiter gentilis* | 13.08.2020 | BE |
| 37 | 210/20 | lib04741 |  | Northern goshawk | *Accipiter gentilis* | 01.08.2020 | BE |
| 38 | 238/20 | lib04742 |  | Northern goshawk | *Accipiter gentilis* | 31.08.2020 | BE |
| 39 | 241/20 | lib04743 |  | Northern goshawk | *Accipiter gentilis* | 23.08.2020 | BE |
| 40 | 244/20 | lib04744 |  | Hooded crow | *Corvus corone cornix* | 18.08.2020 | BE |
| 41 | 219/20 | lib04745 |  | Chilean flamingo | *Phoenicopterus chilensis* | 26.08.2020 | ST |
| 42 | 284/20 | lib04746 |  | Swift parrot | *Lathamus discolor* | 21.09.2020 | ST |
| 43 | 286/20 | lib04747 |  | Horse | *Equus caballus* | 29.09.2020 | ST |
| 44 | 246/20 | lib04757 |  | Golden eagle | *Aquila chrysaetos* | (09.09.2020) survived | BB |
| 45 | 201/20 | na<sup>2</sup> |  | European greenfinch | *Carduelis chloris* | 07.07.2020 | BB |
| 46 | 60/21 | lib04758 |  | Blue tit | *Cyanistes caeruleus* | 2020 | TH |
| C4 | 115/19 | lib04565 |  | Chinese merganser | *Mergus squamatus* | 08.08.2019 | BE |
| C5 | 127/18 | lib04748 |  | Horse | *Equus caballus* | 11.09.2018 | BB |

<sup>1</sup> Abbreviations for federal states: BE, Berlin; BB, Brandenburg; SN, Saxony; ST, Saxony Anhalt; TH, Thuringia
<sup>2</sup> WNV-specific multiplex PCR was unsuccessful due to low amount of WNV-RNA in the sample (Cq 36).

WGS of WNV was performed as described (Quick et al. 2017) with some modifications. Briefly, RNA was reverse transcribed using the SuperScript™ IV First-Strand Synthesis System (Invitrogen) with random hexamers. The cDNA was subjected to the WNV-specific multiplex PCR described in (Sikkema et al. 2020). Using two different primer mixes (Table S1) and an AccuPrime™ *Taq* DNA Polymerase Kit (Invitrogen), two multiplex PCR reactions were performed. Amplicons were purified with 1.8 volume of Agencourt® AMPure XP beads (Beckman Coulter) and quantified using a NanoDrop™ ND1000 Spectrophotometer (Thermo Fisher Scientific). These two purified and quantified amplicon pools were combined per sample in equal concentration (125 ng each) and the volume adjusted to 130 µl. Fragmentation and library preparation steps were performed according to (Wylezich et al. 2018). Quantified libraries (GeneRead DNA Library L Core Kit; QIAGEN) were sequenced using an Ion Torrent S5 XL instrument with Ion 530 chips and respective reagents (Thermo Fisher Scientific) in 400 bp mode according to the manufacturer’s recommendations.

We verified the PCR-based sequencing using five WNV-positive samples from previous seasons (C1-C5; Table S2) that had already been sequenced according to the validated approach described in (Wylezich et al. 2018). Two previously completed libraries of C4 and C5 were enriched for WNV using MyBaits (Wylezich et al. 2018; Wylezich et al. 2021) but still only yielded partial genome sequences. On the contrary, the multiplex PCR-based approach generated complete coding sequences of all 5 test samples, albeit with a truncated 3’end (23-71 nucleotides). The sequences from both approaches were 100% identical for samples C1-C3 and showed a few differences for samples C4 and C5 (Table S2). These results demonstrated that the multiplex PCR approach is suitable for reliable and sensitive WGS of WNV, even from samples with low WNV concentration (up to Cq value 31.5).

Sample #26 (ED-I-258/20) had a genome region with a sequencing depth lower than 30, therefore sequencing results were confirmed with Sanger sequencing. Briefly, cDNA from sample ED-I-258/20 was amplified using additional single-plex PCR assays (primer pairs: WNVUS1_30_LEFT and WNVUS1_30_RIGHT_2, WNVUS1_30_LEFT_2 and WNVUS1_30_RIGHT). The amplicon was sequenced with a BigDye Terminator v1.1 Cycle Sequencing kit (Applied Biosystems™, Thermo Fisher Scientific) on a 3500 Genetic Analyzer Instrument (Applied Biosystems™, Thermo Fisher Scientific).

WNV genome sequences obtained in this study were submitted to the European Nucleotide Archive under the BioProject accession number PRJEB47687.

### 2.5 Datasets

We validated our workflow using two test datasets consisting of WNV complete coding sequences previously characterized and classified into different ranks below the species level. “Test dataset 1” (TD01) consists of 95 WNV whole-genome sequences characterized and classified into different lineages by Fall and colleagues (Fall et al. 2017). Notably, this study considered WNV clades 1a, 1b, 1c, 4a, and 4b/9 as distinct lineages. “Test dataset 2” (TD02) consists of 150 WNV whole-genome sequences allocated to three WNV clades and six WNV clade 1a clusters described by May and colleagues (May et al. 2011). We also combined the sequences from these two test datasets, and a sequence described as a member of the putative WNV clade 1a cluster 7 (Aguilera-Sepulveda et al. 2021). We referred to these sequences as “test dataset 3” (TD03). Available complete coding sequences of WNV lineage 2 and their metadata (e.g. sample collection year and country of origin) were retrieved from GenBank on 10^th^ December, 2021. WNV lineage 2 dataset (WL2) consisted of WNV complete coding sequences from the database and sequences acquired in this study. Accession numbers of WNV sequences per dataset are summarized in Table S3. We also prepared versions of these datasets that excluded sequences with ≥10 ambiguous nucleotides or gaps, and duplicates.

### 2.6 *In-silico* analyses

#### 2.6.1 Sequence assembly

Genome sequences were assembled from raw data using the Roche/454 genome Sequencer software suite v3.0 (Roche). Sequencing adapters and PCR primers were trimmed using the Newbler assembler prior to reference mapping. Initial reference-based mapping against WNV strain 1382/2018/Berlin/Ger (MH986055.1) was done to generate a sample specific consensus sequence. These consensus sequences were then employed as the reference for a second reference-based mapping per dataset. The resulting genome sequences were visually inspected using the Geneious Prime® 2021.0.1 software (Biomatters).

#### 2.6.2 WNV genome characterization and phylogenetic analyses

Complete coding sequences from each dataset (TD01, TD02, TD03, WL2) were aligned using the MUSCLE algorithm (Edgar 2004), and visually inspected using Geneious Prime® 2021.0.1.

#### 2.6.3 Maximum likelihood phylogenetic analysis

The best-fitting nucleotide substitution model for each dataset was calculated using jModelTest 2.1.10 (Darriba et al. 2012). Maximum likelihood (ML) inference with the determined best substitution model and ultrafast bootstrap option (Hoang et al. 2018; Minh et al. 2013) with 100,000 replicates was performed using IQ-TREE 1.6.8 (Nguyen et al. 2015). ML phylogenetic trees were viewed using FigTree software (v1.4.4, http://tree.bio.ed.ac.uk/software/figtree/).

#### 2.6.4 Bayesian phylogenetic analysis

We subjected the dataset consisting of complete genome sequences belonging to the subclade 2.5.3 to the Bayesian Markov Chain Monte Carlo (MCMC) method implemented in the Beast package version 1.10.4 (Drummond and Rambaut 2007; Suchard et al. 2018). We performed regression analyses of the root-to-tip genetic distance in the resulting ML trees against sampling years using TempEst (Rambaut et al. 2016). The spatiotemporal dynamics of WNV and the time to most recent common ancestors (MRCA) were co-estimated using best suited substitution model based on the jModelTesT 2 (Darriba et al. 2012), optimal molecular clock model (relaxed uncorrelated lognormal) and best demographic scenario (the Bayesian SkyGrid coalescent model), which will be explained below.

The optimal molecular clock model (strict or relaxed uncorrelated log normal) and tree prior (Constant, Bayesian GMRF Skyride, or Bayesian Skygrid model) were selected based on the marginal likelihood estimation path sampling and stepping stone sampling methods. The MCMC chain length was run until convergence and sampled every 10^4^ iterations. Convergence was evaluated by approximating the effective sampling size (ESS) after a 10% burn-in using the Tracer software version 1.7.1, with ESS values >> 200 accepted. The strength of the evidence against H0 was evaluated according to Kass and Raftery’s (Kass and Raftery 1995) Bayes factor test as follows: Bayes factor (BF) 1-3 – weak, BF 3-20 – positive, BF 20-150 – Strong, and >150 – very strong (comparison of each parameter summarized in Table S4).

Phylogeographic analysis was performed using a discrete model attributing state characters represented by the detection of location (country) of each strain and the Bayesian stochastic search variable (BSSV) algorithm implemented in BEAST v1.10.4 (Suchard et al. 2018). TreeAnnotator v1.10.4 was employed to summarize the maximum clade credibility (MCC) tree after 10% burn-in and Figtree software v1.4.4 was utilized to visualize the MCC tree. The branches of the trees were color-coded based on the sample’s geographic origin (country).

#### 2.6.5 Affinity propagation clustering (APC)-based workflow for sequence grouping

We analyzed WNV complete coding sequences using a workflow comprising of the APC algorithm and AHC included in the R package “apcluster” (Bodenhofer et al. 2011) implemented in R v4.1.2 and R studio (v2021.09.1-372)(R Core Team 2021). The APC algorithm requires a dissimilarity matrix as input for clustering. For each of the determined clusters, one entity is defined as the “best representative” or the “cluster exemplar”.

Using the Sequence Demarcation Tool (SDT; SDT_Linux64 v1.2) (Muhire et al. 2014), we calculated pairwise global alignments of the coding sequences and from these alignments used the pairwise nucleotide identities to calculate a dissimilarity matrix by subtracting the identities from 1. Subsequently, to increase the robustness and discriminatory power of the APC, these dissimilarities were squared and converted to negative values according to Fischer and colleagues (Fischer et al. 2018) in order to yield the suitable input data for the APC algorithm.

One major problem in clustering is finding the suitable number of clusters to subdivide the dataset into. To this end, Fischer and colleagues (Fischer et al. 2018) developed the “plateau method” to calculate the optimum number of clusters. The number of clusters generated by APC is determined by a parameter called input preference, which by default is set to 0.5. Using the AP clustering algorithm, the suitable “input preference range” from minimum (pmin) to maximum (pmax) can be calculated. For the plateau method, the number of clusters (z-value) is repeatedly determined in dependence of the input preference which is increased in equal steps through the preference range. Usually, with an increase of the input preference, the number of groups monotonously increases; if a reduction occurs, this is deemed a disturbance that leads to the termination of the calculations. Fischer and colleagues defined the best suited number of groups corresponding to the longest plateau that was observed (the same number of clusters observed consecutively for the highest number of iterations before a disturbance occurred). While in principle this was suitable, they nevertheless found that it was not optimal. Since there can be a monotonous increase of the group number without a disturbance occurring throughout the whole preference range, we tested using the last stable plateau as an alternative measure for the definition of the group number. The last stable plateau is defined as the last plateau without disturbance and with at least the set minimal length. For this calculation, we set the minimum number of iterations that make a plateau to 3. Finally, for the definition of the most suitable number of groups present in the input data, the following rules were applied: (i) if both the longest and the last stable plateau resulted in a cluster number higher than the default APC, use the default; (ii) else, if either or both of the plateaus result in values lower than the default, use the higher of the values to set the number of groups. This number of groups was then used to calculate the grouping of the input dataset using the function for AHC from the APC package. The described grouping was applied for the desired number of sub-grouping levels (ranks below the species level). The R code used for these calculations is available as supplemental material.

In order to test the impact of the number of steps and minimum number of iterations to use as the cut-off for definition of the last plateau for the determination of the group number, we used the described test datasets. We ran all calculations with all possible combinations of different step numbers (1,000; 2,000; 5,000; 10,000), minimum plateau lengths (sliding window size 1%, 0.5%, 0.25%, 0.1%, or 0.01% of the step number) and minimum group members to have as input for further sub- grouping (5; 7; 10). In these tests, we found that the coherence of grouping by the described workflow and the phylogenetic trees increased with the number of steps and with the reduction of the sliding window size applied for plateau determination. Notably, with a fixed set of step number and sliding window size, the impact of the minimal group size increases with the increasing size of the input dataset. Since our initial tests showed that ambiguities in the sequences and to a lesser extent also duplicated sequences negatively impact the grouping by the described workflow, we also tested the different test datasets without duplicate sequences and sequences with ≥10 ambiguous nucleotides or gaps. Here, we present results from datasets without sequences having ≥10 ambiguous nucleotides or gaps, and only retained one representative for sequences sharing 100% nucleotide identity. Unless indicated, the used parameter set for the presented results were 10,000 steps, sliding window proportion resulting in sliding window length 3, and minimum group size 5.

#### 2.6.6 Proposal for WNV group designations

Alongside our new workflow, we here propose to use a generic nomenclature based on a hierarchical numbering system. This proposal is outlined in Figure 1. Based on the use of designations in the literature, we propose to designate the levels within the species WNV descending from the species through lineage, clade, subclade, cluster, and finally subcluster. The subclusters can additionally be divided further, then carrying a letter as the suffix. The digits representing the different hierarchical levels are separated by a “.” (compare Figure 1). Here, we examined the grouping in different depths as indicated for the respective analyses. With the lineage designations we followed the established lineage numbering; hence, where necessary, lineage designations automatically assigned in the calculations were replaced by the corresponding established designations.

**Figure 1.**
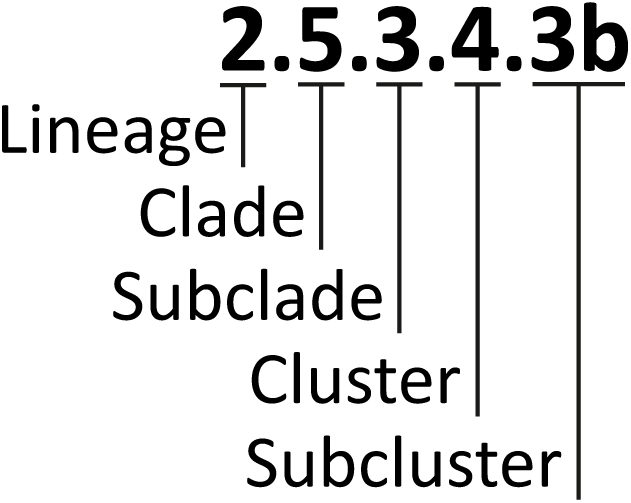
Graphical representation of the proposed hierarchy and the corresponding group labels. The levels of the proposed are ordered top to bottom; the corresponding group label is organized left to right. Note that the subcluster can either have the number only or the number combined with a letter.

#### 2.6.7 Combination of the clustering workflow, phylogenetic analyses and geolocation

The assigned hierarchical levels of WNV sequences detected in Germany from 2018-20 were summarized per new phylogenetic group, collection year, and sample type (wild/captive bird, horse, mosquitoes, and humans). These were exported as a CSV file into the QGIS Desktop (v3.16.15).

## 3. Results and Discussion

Originally, the goal of this study was to provide an update on the WNV epizootic in Germany in 2020. However, we encountered significant problems in consistently allocating WNV sequences into different groups below the species rank, namely:

1) the lack of objective grouping due to undefined demarcation criteria for the splitting of sequences into groups, resulting in arbitrarily adjusted groupings, and
2) the missing common group designations below species level within the West Nile Virus species (and in general) that together with the used nomenclature, which often relies on geographical terms that due to the spread of the virus no longer fit, result in misleading designations.

### 3.1 Proposal for a hierarchical WNV nomenclature below the species level

To date, there is no commonly used system in the WNV research community for the definition and designation of virus groups below the species level. Rather, a substantial number of ways to define and terms to designate virus groups at different levels of a hierarchical system below the species are used. These are also different from what is used for other virus species and what is commonly understood (see Table 2).

**Table 2.** Overview of terms commonly used for the designation of virus sequences into groups below the species rank.

| Term | General definition | Current common use in the WNV research community |  |  | Proposed use |  |
| --- | --- | --- | --- | --- | --- | --- |
|  |  | Definition | Example lineage 1 | Example lineage 2 | Level below species/term | Example designation |
| Lineage | Rank-independent term for the relationships between ancestors and descendants through time (diachronic). Typically, a higher resolution classification compared to clade. (Campbell et al. 2022; Cellinese et al. 2012; Rambaut et al. 2020) | Broadest monophyletic group below the WNV species rank. There are 9 proposed WNV lineages. (Fall et al. 2017) | Lineage 1; Lineage 1a | Lineage 2 | 1 / Lineage | Lineage 1; Lineage 2 |
| Clade | Rank-independent term for a monophyletic group on a phylogenetic tree. Mishler (2010), describe it as "a monophyletic group is all and only the descendants of common ancestors" (synchronic). (Campbell et al. 2022; Cellinese et al. 2012; Rambaut et al. 2020) | Smaller monophyletic group within the lineage. Typically denoted with letters. Example, 1a - 1c and 2a-2d (May et al. 2011; McMullen et al. 2013). In Lineage 2, this level also describes a monophyletic group sharing similar geographic range. (Ziegler et al. 2020) | Clade 1a | Clade 2d; Central/Southern European clade | 2 / Clade | Clade 1.1; Clade 2.5 |
| Subclade | A smaller monophyletic group within a larger clade (Campbell et al. 2022) | Smaller monophyletic group within the clade. More often used in Lineage 2. In Lineage 2, these are also used to describe sequences from a monophyletic branch sharing geographic range. (Barzon et al. 2015; Ziegler et al. 2020) | Not commonly used in Lineage 1 | Central/Southern European subclade; subclade: "Eastern German clade" | 3 / Subclade | Subclade 1.1.4; Subclade 2.5.1 |
|  |  | Definition | Example lineage 1 | Example lineage 2 | Level below species/term | Example designation |
| Cluster | Closely related sequences sharing a certain threshold of nucleotide or amino acid identities, characteristics, or provides to define cluster of disease transmission. (Han et al. 2019) | Smaller monophyletic group within the clade in Lineage 1, sharing a single ancestor and or a fixed unique non-synonymous mutation (May et al). Smaller monophyletic group within the clade or subclade in Lineage 2. (Barzon et al. 2015) | Cluster 1 | Italian Lombardy cluster | 4 / Cluster | Cluster 1.1.4.1 or Subclade 1.1.4 cluster 1; Cluster 2.5.1.1 or Subclade 2.5.1 cluster 1 |
| Subtype | A subset of a species based on a certain characteristic | Below Cluster level, to designate WNV cluster 2 based on geographic location. (May et al. 2011) | Mediterranean subtype (cluster 2) | not commonly used in Lineage 2 | 5 / Subcluster (designated by 5<sup>th</sup> decimal place) | Subcluster 1.1.4.1.6; Subcluster 2.5.1.1.5 |
| Genotype | monophyletic cluster of sequences with high statistical support (Goya et al. 2020) | Used to describe different sequences of WNV lineage 1 cluster 4 detected in America, which shared fixed nonsynonymous mutation. Sub-type was not described in cluster 4 (Mann et al. 2013). | NY99 genotype (cluster 4) | not commonly used in Lineage 2 | 6 / Subcluster (designated by a letter as suffix) | Subcluster 1.1.4.1.6a; Subcluster 2.5.1.1.5a |

The designations of the hierarchical levels *inter alia* include the terms “lineage”, “clade”, and “cluster” (Figure 2). However, the use of the labels to designate different levels of the hierarchical system is variable. The WNV research community especially uses the term “lineage” to describe a broader hierarchical group consisting of clades and/or subclades, while in other virus species, such as severe acute respiratory syndrome coronavirus 2 (SARS-CoV-2) and rabies virus (RABV), the term “clade” defines a broader monophyletic group consisting of subclades and lineages (Campbell et al. 2022; Rambaut et al. 2020).

**Figure 2.**
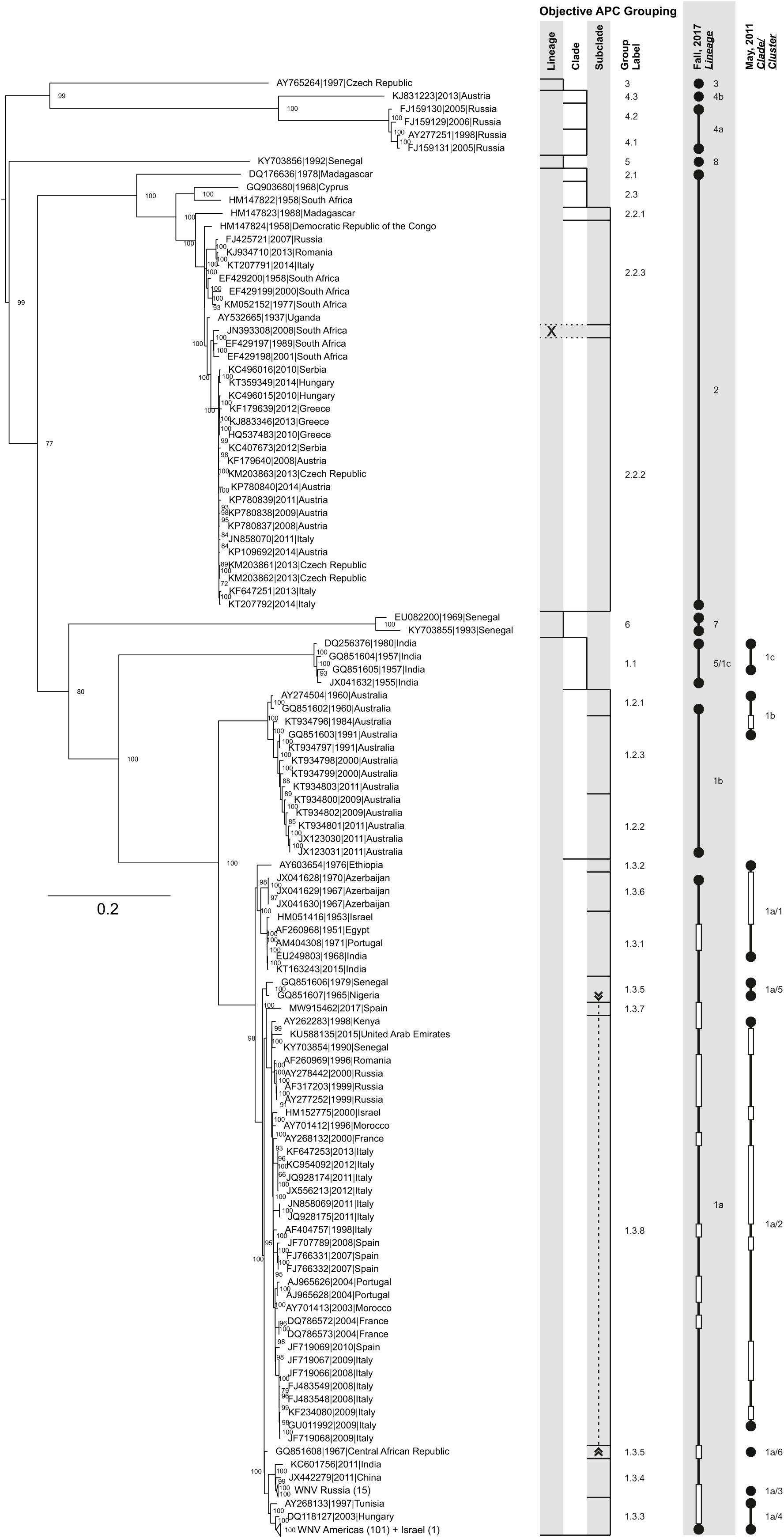
Comparison of APC groupings of test dataset TD03 with previously defined groupings and phylogenetic reconstruction. The representation of the objective APC grouping includes the addressed hierarchical levels, starting with lineage, decreasing from left to right down to the subclade. The vertical lines mark the final level down to which the grouping could be done (limited either by the minimum group size applied for the input of subgrouping or by the hierarchical level that was the last to be shown). Horizontal lines separate the individual groups. Each group is labelled at the right-hand side of the graph. Dashed vertical lines with arrows connect areas of the graph together forming one common group interspersed by other group(s). The horizontal grey rectangle labelled “X” marks a sequence that was not considered for APC/AHC grouping due to its high number of ambiguities (>=10). For comparison, the groupings that were previously published by Fall et al. (2017) and May et al. (2011) are included. Here, a filled circle represents a singleton sequence making up the respective group as labelled and two filled circles connected by a vertical line represent a larger group. White rectangles mark sequences included in the tree but not part of the cited analyses. The maximum likelihood (ML) phylogenetic analysis of sequences from TD03 was done with the best fitting model GTR+I+G and 100,000 ultrafast bootstraps. Few large branches consisting of sequences from almost the same geographic regions are collapsed into triangles. The nodes are labelled with ultrafast bootstrap values.

Moreover, beside the variable use of terms for the designation of hierarchical levels, the criteria used to define the groups are variable. For instance, Aguilera-Sepulveda et al. (2021); Barzon et al. (2015) and May et al. (2011) defined clusters found within WNV clade 1a as sequences belonging to a monophyletic group with a close phylogenetic relationship, with a common ancestor and fixed and unique amino acid substitutions. In another example, McMullen et al. (2013) defined four clades (clades 2a – 2d) based on nucleotide identities and monophyletic branching within the phylogenetic tree. However, the demarcation criteria regarding nucleotide identities or amino acid similarities for each clade were not clearly defined.

Likewise, the labels used to designate the groups are diverse. Often, groups are labelled according to their first geographic occurrence. Although geographic labels may provide epidemiological information regarding the origin of the WNV cases, these descriptive labels can cause misrepresentation. For instance, the geographic range of WNV cases designated to the Lombardy cluster, which consisted of WNV cases from Lombardy, Italy, as of 2015 (Barzon et al. 2015), is recently expanding. The Lombardy cluster now also includes WNV sequences from France and Spain (Aguilera-Sepulveda et al. 2022). Similarly, WNV clade 2d sequences from the European continent were designated according to the supposed region of the viruses’ origins, like WNV sequences from Russia and Romania that were designated as the Eastern European lineage 2 WNV (EE, Figure 3) (Cotar et al. 2018; Ravagnan et al. 2015) or WNV sequences from Hungary, Austria, Greece, Serbia, and Italy that were put into the Central/Southern European lineage 2 WNV (C/SE, Figure 3) (Chaintoutis et al. 2019; Ziegler et al. 2020).

**Figure 3.**
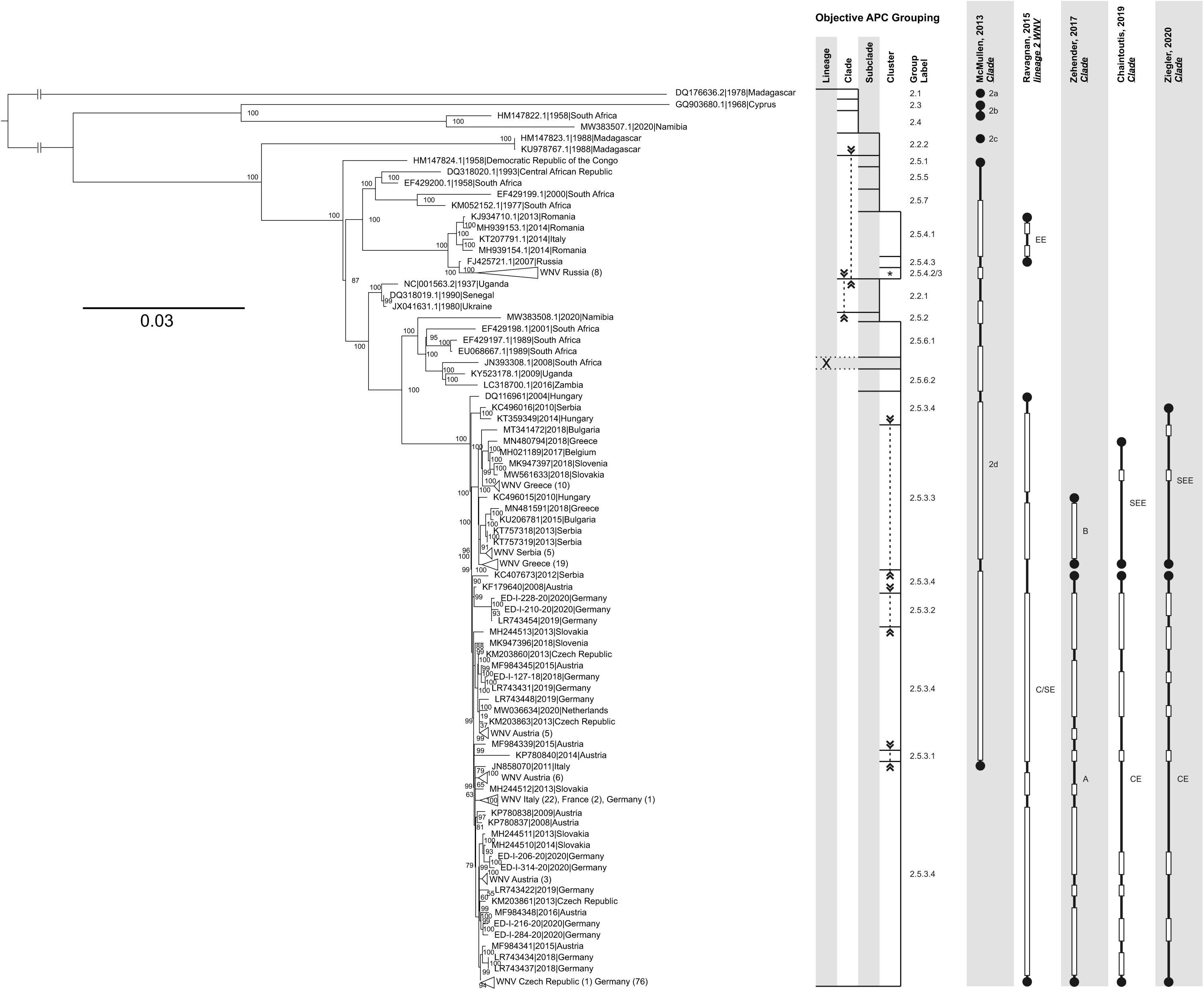
Comparison of APC groupings of WNV lineage 2 (WL2) sequences with previously defined groupings and phylogenetic reconstruction. The representation of the objective APC grouping includes the addressed hierarchical levels, starting with lineage (not calculated here but WNV lineage 2 sequences included according to published references), decreasing from left to right down to the cluster. The vertical lines mark the final level down to which the grouping could be done (limited either by the minimum group size applied for the input of subgrouping or by the hierarchical level that was the last to be shown). Horizontal lines separate the individual groups. Each group is labelled at the right-hand side of the graph. Dashed vertical lines with arrows connect areas of the graph together forming one common group interspersed by other group(s). The horizontal grey rectangle labelled “X” marks a sequence that was not considered for APC/AHC grouping due to its high number of ambiguities (>=10). For comparison, the groupings that were previously published by Chaintoutis et al. (2019), McMullen et al. (2013), Ravagnan et al. (2015), Zehender et al. (2017) and Ziegler et al. (2020) are included. Here, a filled circle represents a singleton sequence making up the respective group as labelled and two filled circles connected by a vertical line represent a larger group. White rectangles mark sequences included in the tree but not part of the cited analyses. The maximum likelihood (ML) phylogenetic analysis of sequences from WL2 was done with the best fitting model GTR+I+G and 100,000 ultrafast bootstraps. Few large branches consisting of sequences from almost the same geographic regions are collapsed into triangles. The nodes are labelled with ultrafast bootstrap values.

Due to the issues outlined above, we set out to design a novel unified system for the hierarchical organization of WNV (and other viruses) based on (I) an objective definition of subgroups (see paragraphs 2.6.5 and 3.2), (II) a defined set of names for the different nested hierarchical levels, and (III) a system for group designations that does not rely on geographic or other names that can likely be subject to change. Although we acknowledge the importance of a universal designation below the species rank encompassing all virus species, we in part still followed the conventional designation of WNV sequences below the species rank to prevent any confusion. For the species *West Nile virus*, we define a term associated with a specific hierarchical level, as summarized in Table 2. We propose to use the following order of hierarchical groups based on increasing shared genetic identities within the group: lineages (highest level below the species, as commonly used in the WNV community, level 1), clade (level 2), subclade (level 3), cluster (level 4), and subcluster (level ≥ 5). Moreover, we propose to utilize a generic nomenclature for the defined groups based on a hierarchical numbering system to designate each group at different hierarchical ranks in a logical and standard manner (Table 2, column “Suggested Usage”). These generic labels also provide information regarding the hierarchical level through the number of decimal and/or alphabetical places included (compare Figure 1). Furthermore, these generic labels can be used continuously even when the group members do not share particular characteristics, such as geographic origin. Finally, we applied these proposals to WNV sequences from previously published studies and members of WNV lineage 2 available in the public database to compare our results with previous classifications.

### 3.2 Application of the developed grouping workflow yields reasonable groups

To address the grouping issues outlined above, we developed a workflow for objective clustering of sequences into different hierarchical groups below species level. This clustering workflow employs APC, which is a non-hierarchical mathematical clustering method, with AHC to split the dataset into groups. This workflow is based on the works of Fischer and colleagues (Fischer et al. 2018), who initially utilized APC to define objective clusters of RABV sequences. Their group also developed the plateau method to determine the number of clusters in a given dataset, typically a user-defined parameter required in clustering programs such as HierBAPS (Cheng et al. 2013), Cluster Picker (Ragonnet-Cronin et al. 2013), TreeCluster (Balaban et al. 2019), and PhyClip (Han et al. 2019). Furthermore, the workflow of Fischer and colleagues only requires pairwise identities between all pairs of virus sequences as input.

Overall, the method overcomes the need for inputting subjective criteria like number of clusters, the minimum number of sequences per cluster, or support thresholds for cluster allocation. While Fischer and colleagues successfully assigned RABV and *Francisella tularensis* isolates into reasonable objectively defined clusters (Busch et al. 2020; Fischer et al. 2018), the APC results were partly incongruent with the branching of a RABV phylogenetic tree. This incongruence is potentially caused by the non-hierarchical clustering properties of the APC algorithm in contrast to the phylogenetic analysis (Fischer et al. 2018), but could also be caused by an uncertainty of the suitable number of clusters present in the dataset. Therefore, to improve the workflow, we further developed the determination of the number of clusters and included AHC to determine the generated clusters. In order to define multiple hierarchical levels, the method was iteratively applied to subsets of the data corresponding to the subgroups of the preceding iteration, i.e. higher level in the hierarchy. For optimization of the parameters, we repetitively analyzed the described test datasets and compared the results with the grouping as described in the respective studies (Chaintoutis et al. 2019; McMullen et al. 2013; Ravagnan et al. 2015; Zehender et al. 2017; Ziegler et al. 2020). We found that the minimum number of sequences per group to be used as input for further subgrouping and the number of iterations used to define the plateau (the window size) had the major impact on the results. On the contrary, the overall number of iterations applied to determine the number of clusters only had less influence. The optimal parameters used for all subsequent analyses were the window size of 3 for the determination of the longest and last stable plateaus, respectively, and the group size of 5 that was necessary to further split the group. In order to ensure that the number of iterations did not limit the quality of the clustering, we used 10,000 iterations throughout.

We initially applied the developed workflow with the settings outlined in the previous paragraph to the test dataset TD03 for the definition of groups within the three proposed levels “lineage”, “clade”, and “subclade”. Figure 2 shows the result of grouping TD03. According to the used minimum size of a group to be used as input for further subdivision in the next lower hierarchical level, the grouping stopped at different levels of the hierarchy. Overall, the objective APC grouping coincides with groups that would be defined when analyzing the tree visually. Most groups we found fit with the traditional definition of a phylogenetic group being monophyletic. In case of the grouping result for TD03, however, we received one subclade (1.3.5) that was not intuitively clear at the first glance at the tree because it was not monophyletic (Figure 2; subclade 1.3.5). This subclade is split in two parts (interspersed by subclades 1.3.7 and 1.3.8), which are in the graph connected with a dashed line with arrows pointing inwards. This split is possible since our workflow mainly depends on the nucleotide identities of pairwise aligned sequences but not on reconstructed hierarchical connections. Looking at the tree in more detail, it becomes clear that the branch lengths between the three subclade members are indeed quite short and therefore the grouping makes sense. Hence, we proceeded with the proof-of-concept for the developed method.

### 3.3 Proof-of-concept for the developed clustering workflow

For the proof-of-concept, we compared our grouping results with published groupings. Using the abovementioned parameters, we could reproduce the groupings of test datasets TD01 and TD02 as published (Fall et al. 2017; May et al. 2011) (results not shown). For the combined dataset TD03, we obtained the grouping shown in Figure 2. Both Fall and colleagues and Rizzoli and colleagues categorized WNV lineages 1a, 1b, 1c, 4a, and 4b (4/9) as distinct and separate lineages (Fall et al. 2017) (Rizzoli et al. 2015), while May and colleagues designated the same groups of sequences belonging as clades 1a, 1b, and 1c which they further subdivided into clusters (May et al. 2011). As can be seen, the objective grouping of the APC/AHC workflow overall coincides with the previously performed groupings, albeit at different levels of the hierarchy and hence different labels. At the lineage level, although lineages 1a, 1b, and 1c (Fall) and 4a and 4b (Fall), respectively, are fused together in one group each by the APC/AHC workflow, the new defined lineages match those of Fall and colleagues (Fall et al. 2017). At the next level (clade), our workflow divides the fused lineages into clades, with lineage 4b (Fall et al. 2017) corresponding to clade 4.3 and lineage 4a (Fall et al. 2017) being subdivided into clades 4.1 and 4.2. Likewise, lineages 1a, 1b, and 1c (Fall et al. 2017) correspond to clades 1.1 (1c), 1.2 (1b), and 1.3 (1a). At the subclade level of our proposed nomenclature, the clusters that May et al. (2011) defined within lineage 1a match our subclades quite well: the members of the cluster 1a/1 (May et al. 2011) are comprised within the subclades 1.3.1 and 1.3.2; sequences of cluster 1a/2 are comprised in subclade 1.3.8; 1a/3 and 1a/4 correspond with subclades 1.3.4 and 1.3.3, respectively; finally, clusters 1a/5 and 1a/6 are combined into subclade 1.3.5. Notably, subclade 1.3.5 is not a monophyletic group but in the phylogenetic tree all descend from the same branch and their branch lengths are very short. Therefore, the co-allocation by APC/AHC is congruent with the minor distances that are visible in the phylogenetic tree. In summary, although in detail there are a few differences, overall, the developed objective grouping by APC/AHC yields meaningful and reliable groupings.

In addition to the above proof of concept study for the separation of WNV of all lineages into the different hierarchical levels (lineages, clades, and subclades), we analyzed the WNV lineage 2 complete coding sequences available in the INSDC databases. As stated above for the first analysis, the grouping we received overall fit well with what is seen intuitively in the tree. Usually, the observed polyphyletic interspersed groups, e.g., clades 2.2 and 2.5 in Figure 3, which are in part associated with low ultrafast bootstrap values in the tree (according to the IQ-Tree documentation, only values above 95 % indicate trustworthy clades (Minh et al. 2022)) are resolved at the next lower grouping level (in this example at the subclade level). Here, clade 2.2 (Figure 3) is a polyphyletic group comprising five sequences, which are at the subclade level separated into subclades 2.2.1 and 2.2.2. This interspersed grouping at the clade level, which occurs in the APC step based on the pairwise identities, cannot be resolved using AHC. This incongruency is due to the inherent non-hierarchical characteristics of the APC, as described by Fischer and colleagues (2018). Similarly, in the deeper grouping of subclade 2.5.3 sequences, subcluster 2.5.3.4.3a includes WNV sequences that are interspersed in the ML and MCC trees (Figure 4). This subcluster formed a paraphyletic group in both ML and MCC trees, and demonstrated low ultrafast bootstrap values (<80%) and posterior probability values (<0.6), respectively.

**Figure 4.**
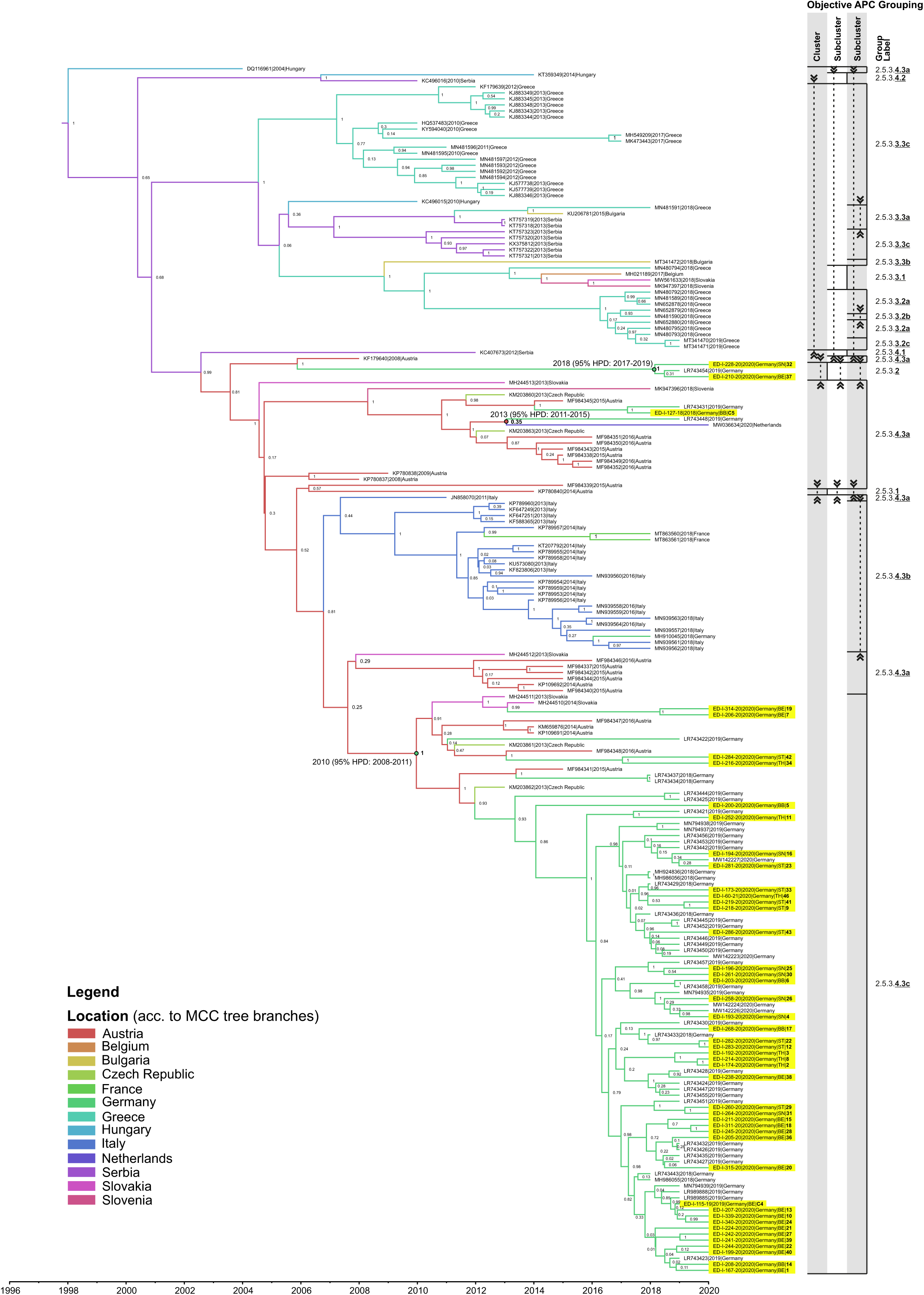
Bayesian maximum clade credibility (MCC) tree representing time scaled phylogeny of European WNV subclade 2.5.3 complete coding sequences together with objective APC groups. WNV sequences acquired in this study are highlighted yellow. All other WNV sequences were retrieved from GenBank and are listed in Table S3. The colored branches of MCC trees represent the most probable geographic location of their descendants (see legend “locations”). Bayesian posterior probabilities are indicated at each node. Time (in years) is indicated as x-axis below the MCC tree. The time for the most recent common ancestor (MRCA), time intervals defined by the 95% highest posterior density (95% HPD), and posterior probabilities (pp) are shown in the following nodes that consist of the following WNV sequences: (i) LR743448 and MW036634, (ii) cluster 2.5.3.2 sequences, and (iii) subcluster 2.5.3.4.3c sequences. The representation of the objective APC grouping includes the addressed hierarchical levels, starting with cluster decreasing from left to right down to the subcluster. The vertical lines mark the final level down to which the grouping could be done (limited either by the minimum group size applied for the input of subgrouping or by the hierarchical level that was the last to be shown). Horizontal lines separate the individual groups. Each group is labelled at the right-hand side of the graph. Dashed vertical lines with arrows connect areas of the graph together forming one common group interspersed by other group(s).

The discussed topology in phylogenetic trees depicts the so-called “supercluster”, wherein divergent subgroups are nested within a more extensive cluster structure (Han et al. 2019). Therefore, in combination with phylogenetic trees our grouping workflow can also provide insights regarding the source-sink ecological dynamics of WNV lineage 2 in Europe. This dynamic has been previously discussed in the phylogeographic and phylodynamic analyses of Zehender et al. (2017) and Ziegler et al. (2020). Specifically, cluster 2.5.3.4 may represent the putative source of the WNV population that gives rise to its subgroups, reflecting the trajectory and divergence of variants (Han et al. 2019). In parallel, members of cluster 2.5.3.4 were detected in locations described as “radiation centers or sources” of WNV lineage 2 migration in Europe (e.g., Hungary and Austria). Furthermore, members of other WNV clusters were detected in areas described as “receiving areas or sinks” of WNV migration, such as Greece (cluster 2.5.3.3).

To further verify the workflow, we compared our grouping with previously published results of McMullen et al. (2013), Ravagnan et al. (2015), Zehender et al. (2017), Chaintoutis et al. (2019) and Ziegler et al. (2020). Noteworthy, all studies that were available for comparison only included partial sets of the sequences that we included here. The comparison of the results of the objective APC/AHC grouping and the clades defined by McMullen (McMullen et al. 2013) shows that there are two main differences between both: (i) McMullen’s clade 2b is disrupted into clades 2.3 and 2.4 in our grouping; this is likely caused by the inclusion of the 2020 sequence from Namibia (MW383507), which forms clade 2.4 together with the 1958 South African sequence (HM147822) that was included in McMullen’s clade 2b; (ii) the sequences comprised in McMullen’s clade 2d were now put into clade 2.5, except for 1990 Senegal (DQ318019) and 1937 Uganda (NC_001563) that form clade 2.2 together with one 1988 sequence from Madagascar (HM147823). These two deviations show the expectable effect of addition of sequences on tree topology and sequence grouping. The comparison between the groupings of Ravagnan and colleagues (Ravagnan et al. 2015) and ours shows that the virus group designated “Eastern European lineage 2 WNV” (labelled EE in Figure 3) coincides with our subclade 2.5.4 and those of the “Central/Southern European lineage 2 WNV” (labelled C/SE in Figure 3) are all grouped into subclade 2.5.3. In the studies of Zehender et al. (2017), Chaintoutis et al. (2019) and Ziegler et al. (2020), viruses belonging to Ravagnan’s C/SE lineage 2 WNV (Ravagnan et al. 2015) were subdivided into two groups. These were labelled clade A (Zehender) or Central and Eastern European clade (CEC; Ziegler; Chaintoutis) and clade B (Zehender) or Southeastern European clade (SEEC; Ziegler; Chaintoutis), respectively. Using our APC/AHC workflow, they are grouped together in subclade 2.5.3. At the next hierarchical level (cluster), with a single exception (LR743454, Germany 2019, cluster 2.5.3.2), clade A/CEC is completely comprised within cluster 2.5.3.4. Likewise, clade B / SEEC is fully comprised in cluster 2.5.3.3, except for the two sequences from Hungary 2014 (KT359349) and Serbia 2010 (KC496016). Interestingly, cluster 2.5.3.1 comprises only a single WNV sequence from Austria (KP780840) that has not been included in previous phylogenetic studies (Chaintoutis et al. 2019; Ziegler et al. 2020) since it was considered an outlier based on its temporal signal relative to other WNV subclade 2.5.3 sequences. This sequence also showed the lowest pairwise nucleotide identities among members of subclade 2.5.3. Noteworthy, Ziegler and colleagues highlighted that LR743454 formed its own distinct subclade within the CEC. In our analysis, this sequence received two companions, altogether forming cluster 2.5.3.2.

Taken together, the presented comparisons between published studies and the grouping obtained by application of the newly developed APC/AHC workflow show that our objective workflow reliably puts sequences into meaningful groups.

### 3.4 WNV circulation in Germany extended in space and species

In 2020, we detected 65 birds (captive = 33 and wild = 32) and 22 horses that tested positive for WNV in Germany (diagnosed between July 14 and October 20 and two retrospective cases from 2021). All but one WNV-positive bird succumbed to infection (Table 1; #44). The number of notifiable cases of WNV in birds and horses in 2020 is similar to the previous year, particularly in regions with the highest WNV activity i.e., Berlin, Saxony, and Saxony-Anhalt (Figure 5) (Ziegler et al. 2020). However, we observed an increasing number of WNV-cases in Brandenburg, Thuringia, and Lower Saxony. All WNV-positive birds and horses detected in 2020 were found in federal states which also reported WNV cases in 2018 and 2019 (Figures 5 and S1) except for a new WNV-case detected in Lower Saxony. Notably, all 22 probable autochthonous human WNV cases in 2020 occurred in these federal states (Berlin = 7; Saxony = 11; Saxony Anhalt = 4) (European Centre for Disease Prevention and Control 2020; Frank et al. 2022; Pietsch et al. 2020). Therefore, this kind of WNV surveillance in both wildlife and captive animals could provide an early warning for autochthonous WNV-infection in humans in Germany. Hence, reports of WNV infection in birds and horses in an area must be provided promptly (e.g., updates of FLI websites) to advise the medical community and the public regarding a potential risk of WNV-infection in specific regions in Germany, as well as the risks in blood transfusion and organ transplantation safety. Although vaccines against WNV disease in humans are still under development (Ulbert 2019), clinicians must be aware of the potential presence of WNV circulation in the local region to reach a correct diagnosis since WNV diagnostics is not routinely performed in Germany (Schneider et al. 2021).

**Figure 5.**
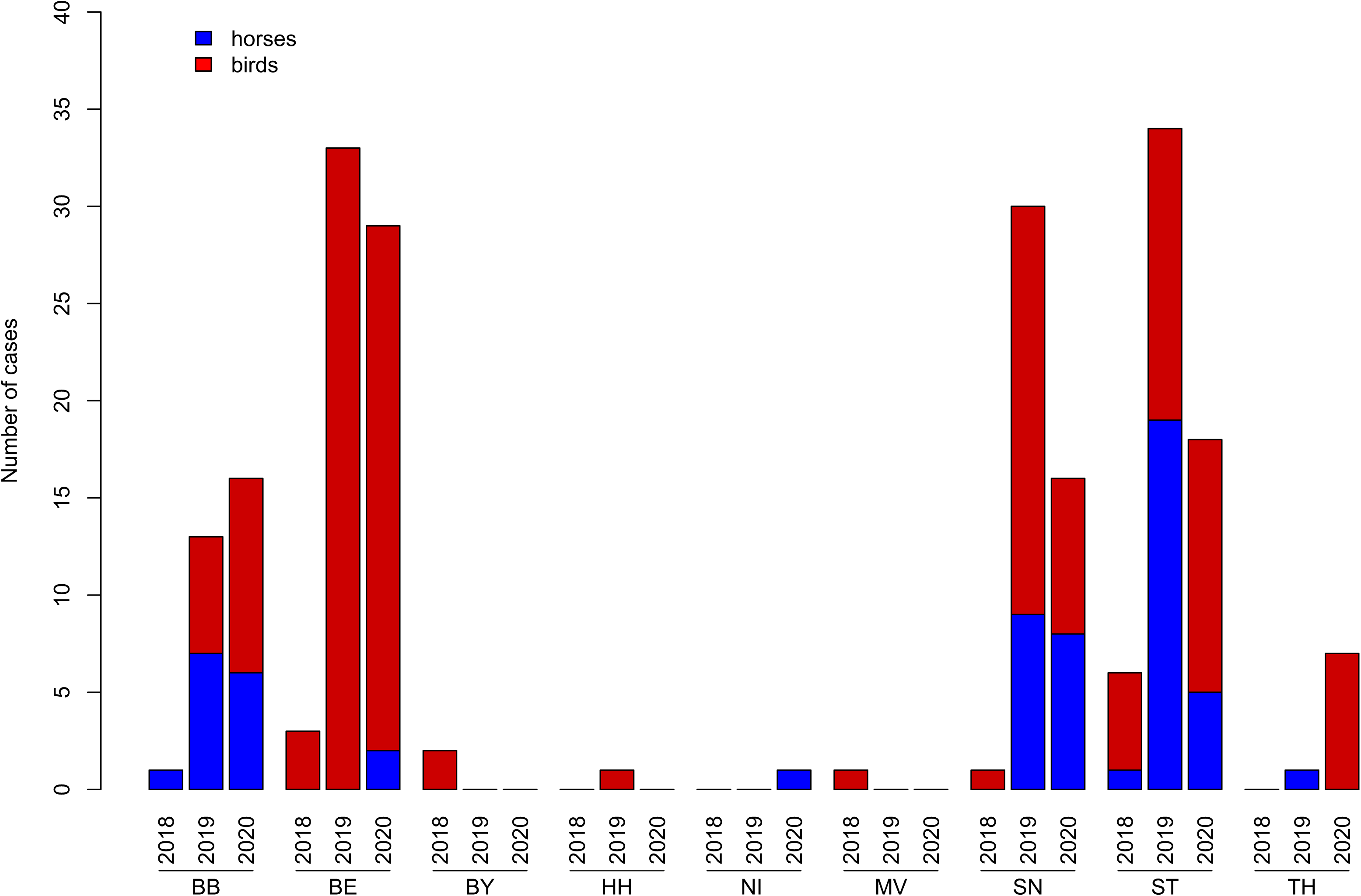
Notifiable WNV-cases of birds and horses in Germany from 2018 –20. The number of cases were summed up per federal state and year. Notifiable cases in horses and birds were represented by blue and red bars, respectively. Abbreviations of federal states in Germany: BB – Brandenburg, BE – Berlin, BY – Bavaria, HH – Hamburg, NI – Lower Saxony, MV - Mecklenburg Western Pomerania, SN – Saxony, ST – Saxony-Anhalt, and TH – Thuringia.

Here we report the first case of a WNV-infection in Lower Saxony, where a horse with WNV-specific IgM antibodies was detected in the Helmstedt district, and the first reported cases of WNV-infected birds in Thuringia, particularly in the districts of Erfurt and Gera (Figures 5 and S1). We also reported the first cases of WNV-infection in three districts in Brandenburg (i.e., Teltow-Fläming, Barnim and Dahme-Spreewald) and in one district in Saxony-Anhalt (Börde) (Figure S1). Areas with reported WNV-infection match with areas with high average temperatures (>20°C), lower average precipitation (≤250 mm), and lower average climatic water balance (-150 – 50 mm) in summer 2020 (Figure S2) (Deutscher Wetterdienst 2020). Higher average temperatures over several days may increase the risks of WNV transmission through mosquito vectors (Holicki et al. 2020). The higher average temperatures in these areas probably caused the epizootic emergence of WNV by shortening the extrinsic incubation period (EIP) in local mosquito populations. Furthermore, WNV activity is more likely to increase during drought than during rainy periods (Paull et al. 2017). It is also possible that the declining water sources force the avian reservoir hosts to aggregate, increasing the probability of contact between birds and mosquitoes and WNV transmission (Paull et al. 2017). However, we did not detect the re-emergence of WNV in Hamburg, and in two districts in Brandenburg (Ostprignitz-Ruppin and Havelland) in 2020, despite the observed higher average temperatures (>20 °C) and lower average precipitations (126-200 mm) in summer 2020.

We also detected WNV infections in 21 different bird species from six taxonomic orders (Table 3). The majority of WNV-infected avian species are classified as birds of prey (order *Accipitriformes*, 29%), followed by songbirds (order *Passeriformes*, 26%), captive flamingos (order *Phoenicopteriformes*, 23%) and owls (order *Strigiformes*, 17%). Most of the WNV-infected bird species in 2020 were also reported in an earlier study (Ziegler et al. 2020), except for the alpine chough *(Pyrrhocorax graculus*), Bohemian waxwing *(Bombycilla garrulus)*, and golden eagle (*Aquila chrysaetos*). However, all three species belong to taxonomic orders that were already described before to be repeatedly affected by WNV (*Passeriformes* and *Accipitriformes*) (Michel et al. 2018; Michel et al. 2019). Notably, the golden eagle from Brandenburg (#44) is the only reported case in 2020 that recovered from WNV-infection (Table 1).

**Table 3.**
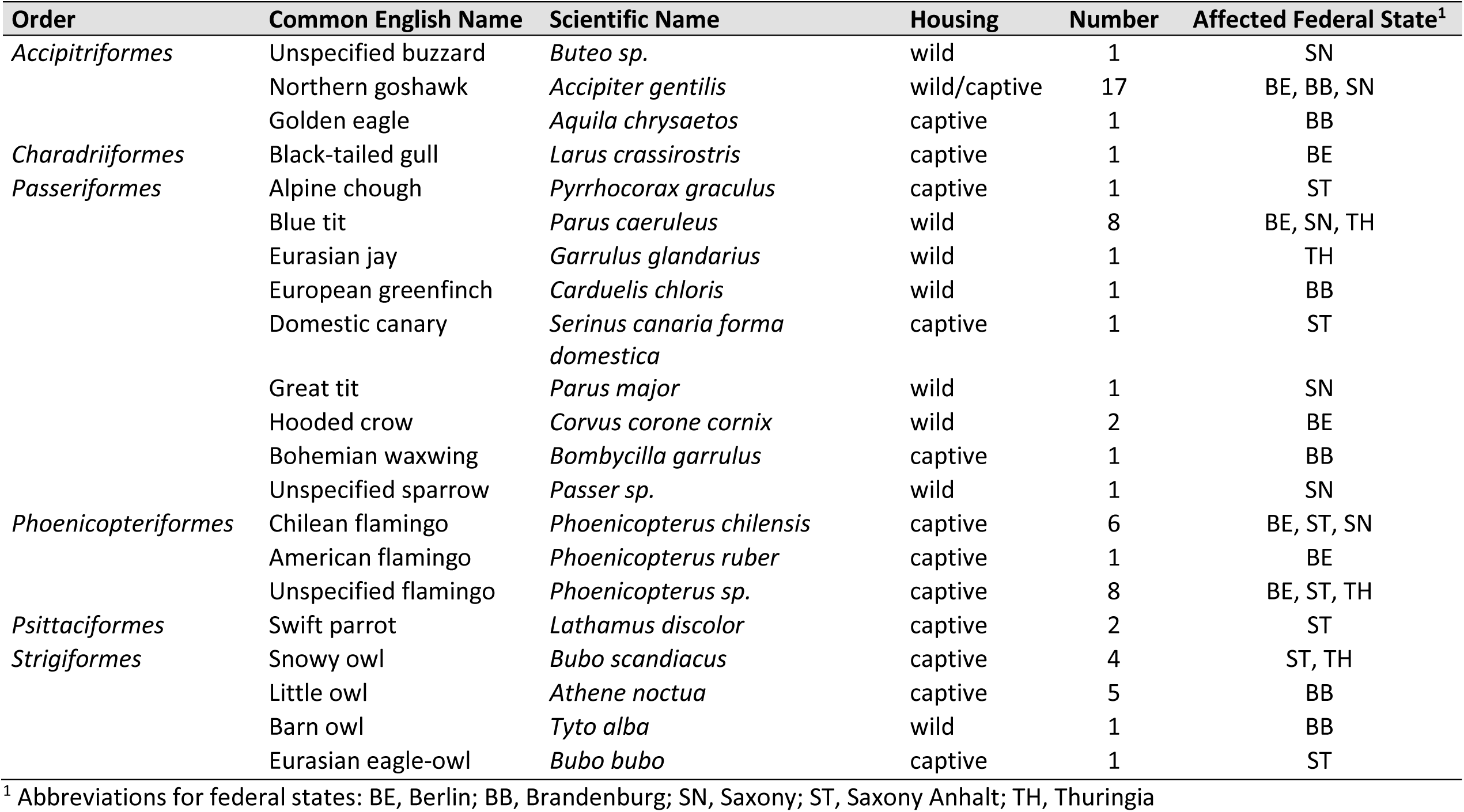
Summary of avian species infected with WNV in 2020 in Germany

### 3.5 Update on the WNV situation in Germany, 2020

After we had validated the workflow, we analyzed the ongoing WNV epizootic in Germany using this tool. The result of grouping sequences that belong to subclade 2.5.3, to which all viruses circulating in Germany until 2020 belong, is shown in Figure 4. As can be seen, subclade 2.5.3 can be further subdivided into the four clusters 2.5.3.1, 2.5.3.2, 2.5.3.3, and 2.5.3.4. Interestingly, cluster 2.5.3.1 only comprises the beforementioned Austrian sequence KP780840 that was previously deemed an outlier and therefore disregarded in previous analyses. Cluster 2.5.3.2, which can due to the group size restriction also not be further subdivided, consists of three German sequences (LR743454 from 2019 and #32 and #37 from 2020), also mentioned above. On the contrary, clusters 2.5.3.3 and 2.5.3.4 can be further subdivided into multiple subclusters each. Although to a large extent the detected subclusters comprise sequences from individual countries, they are clearly not geographically homogenous, highlighting the problem of geographic criteria for the designation of phylogenetic groups. For instance, subcluster 2.5.3.4.3b mainly comprises sequences from Italy but also 2 from France and 1 sequence of a case imported to Germany (MH910045). Likewise, subcluster 2.5.3.4.3c, into which the majority of WNV sequences from Germany were grouped, also comprises sequences from Slovakia (n=2), Austria (n=5), and the Czech Republic (n=2).

As summarized in Figures 6 and 7, sequences from WNV circulating in Germany from 2018-20 were allocated to cluster 2.5.3.2 and subclusters 2.5.3.4.3a and 2.5.3.4.3c, respectively. A sequence of cluster 2.5.3.2 was first detected in 2019 (LR743454) and previously formed an outlier (Ziegler 2020) but now two additional viruses of this cluster were detected (ED-I-228-20 - #32, ED-I-210-20 - #37) (Figure 6). The MRCA of WNV in cluster 2.5.3.2 (see Figure 4) was estimated to have existed around 2018 (95% highest posterior density or 95% HPD: 2017- 2019; Bayesian posterior probabilities or pp: 100%). Unlike viruses of cluster 2.5.3.2, viruses of subcluster 2.5.3.4.3a were only detected in 2018 (ED-I-127-18 – C5) and 2019 (LR743431, LR743448), but not in 2020 (Figure 6). Given the available WNV genome sequences, we cannot confirm whether these minor genotypes (cluster 2.5.3.2 and subcluster 2.5.3.4.3a) have successfully overwintered or been introduced to Germany in separate events. Furthermore, we may have missed these minor WNV clusters and subclusters as we could not sequence all WNV-positive cases from 2018-20 (Table 1; #45). For instance, most horse samples are serologically WNV IgM positive but WNV-RNA negative, preventing the successful sequencing of WNV genomes. Moreover, organ materials from small passerines were often depleted after necessary routine diagnostics at the regional veterinary laboratories for other relevant avian viruses or after confirmatory diagnostics at the national reference laboratory at the FLI. In some cases, simply the sample quality and/or quantity prevents from generating the genome sequences, despite the use of the WNV multiplex-PCR-based HTS approach (Sikkema et al. 2020).

**Figure 6.**
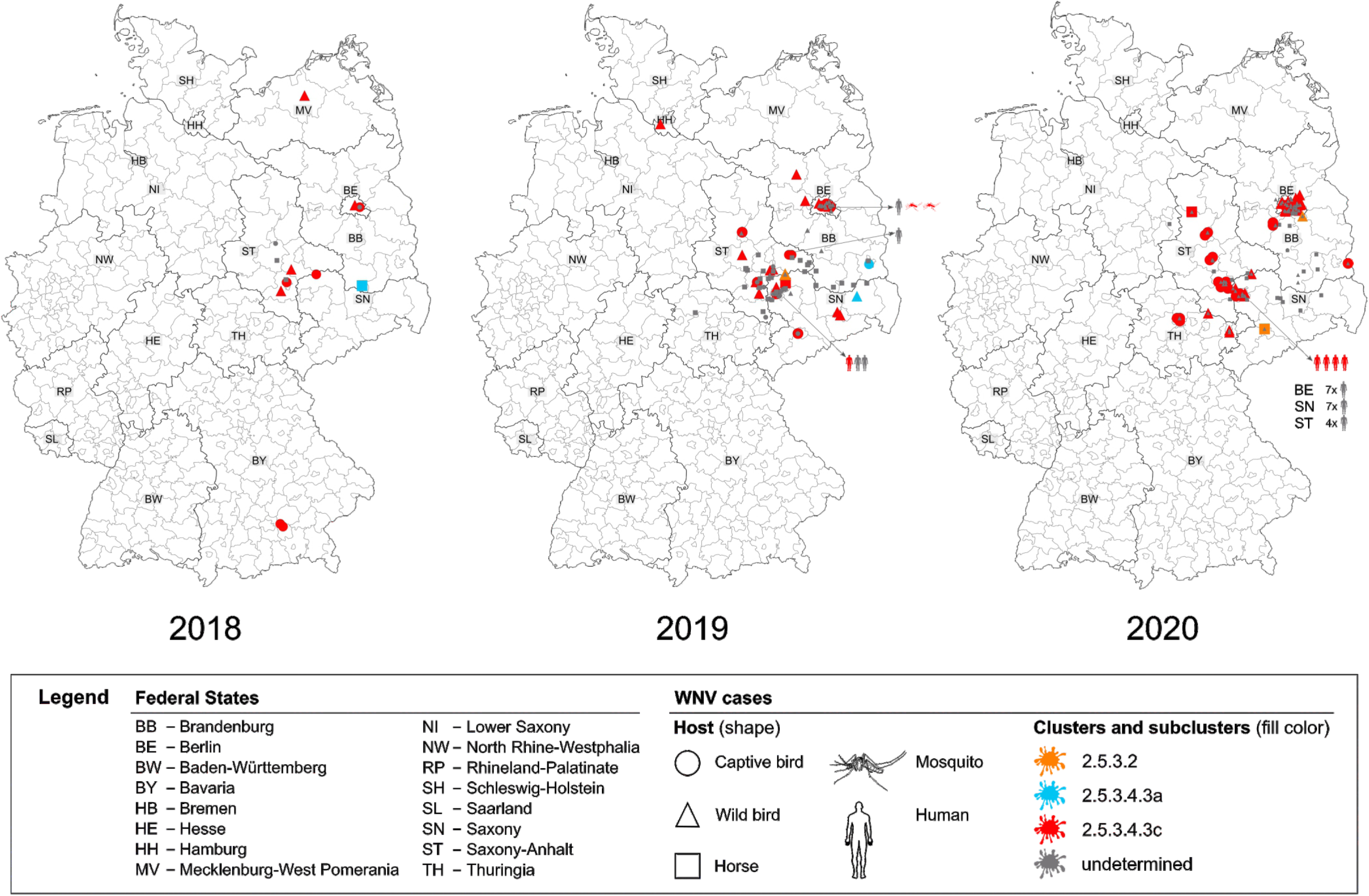
Geographic distribution of WNV cases in Germany from 2018-20 per host and (sub)clusters. Labelling according to the legend in the graph. WNV-positive cases confirmed by the National Reference Laboratory without complete coding sequences are depicted in grey (labelled “undetermined” in the legend).

**Figure 7.**
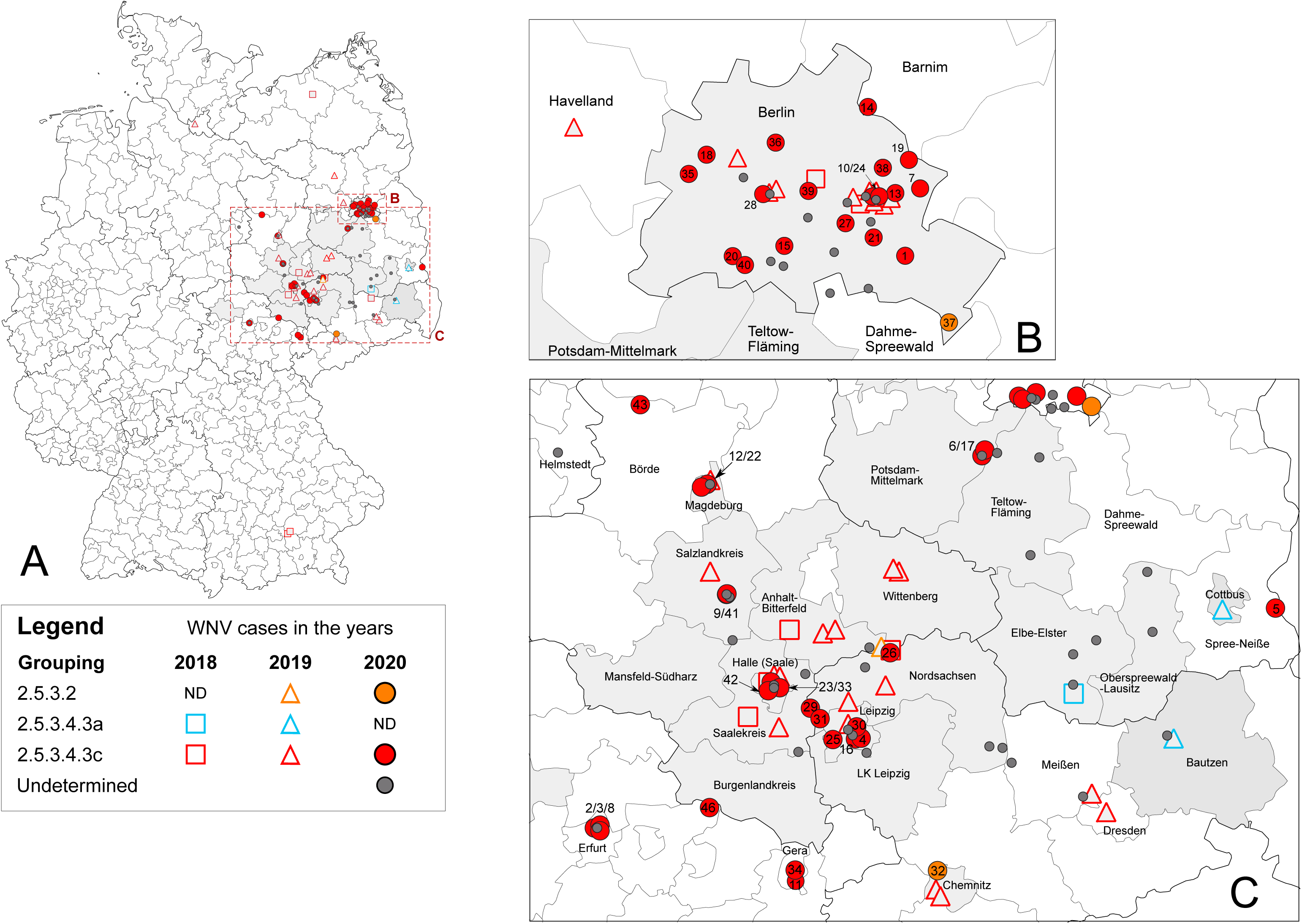
Summarized geographic distribution of WNV cases in Germany from 2018 –20. Labelling according to the legend in the graph. WNV-positive cases confirmed by the National Reference Laboratory without complete coding sequences are depicted in grey (labelled “undetermined” in the legend). Districts colored gray indicate areas with (additional) WNV-positive cases from WNV seasons 2018-19 without a complete coding sequence. Areas with high WNV activity in 2020 are shown in enlarged and separated maps, (B) Berlin, (C) Saxony, Saxony-Anhalt and Thuringia. New WNV cases from this study are indicated with numbers as described in Table 1.

Beside the above mentioned two minor groups, the vast majority of WNV circulating in Germany were allocated to subcluster 2.5.3.4.3c, which comprises all sequences previously allocated to the EGC plus additional sequences, inter alia two previously defined minor subclades comprising sequences LR743422 and LR743437/LR743434 (Ziegler et al. 2020). The EGC, which was the dominant genotype that circulated in Germany from 2018-19, was characterized by a unique non-synonymous mutation (Lys2114Arg) located within the NS3 encoding genome region (noteworthy, LR743444 and LR743425 were previously designated into the EGC but do not harbor this mutation). This mutation no longer is a marker of the respective group (subcluster 2.5.3.4.3c) which also comprises sequences without that specific mutation. Overall, the grouping we now observed (one major subclade and two minor (sub)clusters agrees with our previous WNV report, wherein we detected six distinct “subclades” circulating in Germany in 2018 and 2019 (Ziegler et al. 2020).

We estimated that the MRCA of the monophyletic branch consisting of subcluster 2.5.3.4.3c sequences existed around 2010 (95% HPD: 2008–2011; pp: 100%). Despite the fact that the vast majority of the 2.5.3.4.3c sequences are from Germany, it appears highly unlikely that the ancestors of that subcluster evolved in Germany, as confirmed WNV-positive cases in Germany were only detected from 2018 onwards by the extensive arbovirus monitoring performed in the country since 2011 (Michel et al. 2019; Ziegler et al. 2022). Rather, given that (i) the estimated MRCA of the EGC coincided with large reported outbreaks in eastern and southeastern Europe (Aberle et al. 2018; Jungbauer et al. 2015; Kolodziejek et al. 2015; Kolodziejek et al. 2018; Rudolf et al. 2014; Sedlak et al. 2014; Vlckova et al. 2015) and (ii) WNV complete genomes are not available from neighboring countries, we cannot determine where this subcluster diverged. Therefore, we hypothesize that members of the EGC were more likely introduced to Germany from neighboring countries in separate events and in a later time than its estimated MRCA.

While we detected subcluster 2.5.3.4.3c all over the WNV affected regions in Germany from 2018 until 2020, making it the dominating subcluster, viruses of (sub)cluster 2.5.3.2 and 2.5.3.4.3a were both in time and space restricted and of minor impact for the ongoing epizootic (Figures 6 and 7). Like with the sporadic occurrence of the aforementioned two (sub)clusters, there are also regions within Germany where WNV occurrence is only sporadic (regardless of the virus’ phylogenetic group). Namely, we detected WNV infected wild birds in Rostock, Mecklenburg-Western Pomerania in 2018 (n=1) and in Hamburg (n=1), Havelland, Brandenburg (n=1) in 2019. However, in these areas in the succeeding years, WNV activity was not reported.

As in the preceding years, in 2020, except for two cases in which viruses of cluster 2.5.3.2 were detected, all other viruses were grouped into subcluster 2.5.3.4.3c, and in the same cities and districts as before (Figure 7). In addition, viruses of subcluster 2.5.3.4.3c were detected in three districts in Thuringia. These observations suggest that viruses of subcluster 2.5.3.4.3c successfully established in local avian and mosquito populations in the affected regions, namely in Berlin, Saxony (particularly within Leipzig and neighboring areas) and Saxony-Anhalt, which led to the endemic circulation of WNV in these areas in 2020. We also observed the continuous geographic expansion of WNV belonging to subcluster 2.5.3.4.3c from 2018 to 2020; however, only time will tell whether members of this subcluster successfully overwinter and establish themselves in these newly affected areas. In 2021, however, WNV cases in birds and horses were predominantly reported in Berlin, with a few additional WNV-cases reported in Saxony, Saxony-Anhalt, and Brandenburg (FLI report).

WNV sequences within subcluster 2.5.3.4.3c from Germany were acquired from mosquito pools (n=2), horses (n=2) and different bird species (n=78) belonging to seven taxonomic orders. Complete coding sequences from five human WNV cases reported from 2019 (n=1) to 2020 (n=4) were also allocated into WNV subcluster 2.5.3.4.3c (Figure 6). We excluded a few human WNV cases where either only a partial genome sequence (n=2) (Pietsch et al. 2020; Ziegler et al. 2020) or no sequence information at all (n=3) (Ziegler et al. 2020) was available. These WNV cases did not meet the required criteria for the APC/AHC grouping, i.e., WNV complete coding sequences with <10 nucleotide gaps or ambiguities. As expected, the available partial WNV genome sequences of the two human cases (MN794936, MW142225) had the highest sequence identities with members of the subcluster 2.5.3.4.3c. In addition, recently published complete coding sequences (MZ964751.1, MZ964752.1, MZ964753.1) from three human WNV cases reported in 2021 (Schneider et al. 2021) have the highest sequence identities with members of subcluster 2.5.3.4.3c. Therefore, as of writing, only members of subcluster 2.5.3.4.3c have been reported to cause WNV infection in humans in Germany. Members of subcluster 2.5.3.4.3a, likewise detected in Germany, have previously been reported to cause human WNV- infection in other countries, i.e. Austria (Kolodziejek et al. 2015; Kolodziejek et al. 2018). The higher spread and frequency of subcluster 2.5.3.4.3c in Germany are the likely cause for it being the sole subcluster so far associated with human WNV cases reported in Germany.

Here, we also obtained the complete coding sequence of WNV detected in a horse from 2018 (C5), grouping in subcluster 2.5.3.4.3a (Figures 4 and 7). Viruses of subcluster 2.5.3.4.3a are found widespread across Europe over a long period of time, e.g., in Italy (2011), Austria (2015-2016), the Czech Republic (2013), Slovakia (2013), Slovenia (2018), Germany (2018-2019), and the Netherlands (2020) (Figure 4). Noteworthy, we did not find any member of this subcluster among the sequenced WNV cases in 2020. Still, we cannot directly conclude that its absence in 2020 was due to a failed establishment in Germany since we were not successful in generating sequences from all 65 WNV PCR-positive birds from the 2020 season. The MRCA of WNV MW036634, detected in a Culex mosquito pool collected in Utrecht, the Netherlands, in 2020 (Sikkema et al. 2020) and LR743448 (collected in Cottbus, Brandenburg, Germany in 2019) was predicted to exist around 2013 (HPD 95%: 2011-2015 and pp: 35%) (Figure 4). However, these WNV cases from Cottbus and Utrecht were detected >600 km apart within a short period. Given the large distance between Utrecht and Cottbus together with the ubiquitous distribution of subcluster 2.5.3.4.3a in Europe, we suspect that these two WNV cases might be independent of each other, although they are the closest known relatives. Due to the greater distances between the Netherlands and those regions of Europe where related WNV were previously detected, we hypothesize that different modes of WNV dispersal other than bird migration may have played a role to the WNV introduction in the Netherlands. For instance, the translocation of WNV-infected mosquitoes inside vehicles (planes, ships, automobiles) may have occurred as described for different mosquito species (Bakran-Lebl et al. 2021; Brown et al. 2012; Eritja et al. 2017; Ronca et al. 2021).

## Conclusions

Here, we introduced a structured and unbiased clustering workflow to systematically allocate WNV complete coding sequences to at least six hierarchical groups below the species level: **lineages, clades, subclades, clusters,** and **subclusters**. In addition, we propose a generic hierarchical decimal numbering system designating each group below species rank. We successfully applied the method to allocate WNVs into groups below the species level and this workflow can also be applied to classify other virus species into hierarchical subgroups. Our workflow only requires a matrix of pairwise sequence identities as input. Essential parameters (e.g. number of clusters, threshold, etc.) are entirely decided by the mathematical algorithm, thus removing subjective input from users. Furthermore, the results of our workflow can be combined with different analyses, such as the classical phylogenetic ML tree and the time-scaled MCC tree.

Our analyses revealed that subcluster 2.5.3.4.3c was the predominant WNV subcluster circulating in Germany from 2018-20, accompanied by co-circulating minor WNV (sub)clusters. This finding indicates that the WNV genetic diversity in Germany is primarily influenced by the successful establishment, enzootic maintenance and expansion of subcluster 2.5.3.4.3c, possibly supplemented with continuous incursion and potential overwintering of WNV of other (sub)clusters. These other (sub)clusters detected in Germany overlapped in space and time with the dominant subcluster 2.5.3.4.3c. The minor groups were found in both wild and captive birds, as well as in horses. Therefore, to obtain the full picture of WNV circulation, it will be necessary to obtain whole-genome sequences from all WNV-cases whenever possible, to ensure that also minorities are found.

Since all human WNV cases in 2020 occurred in WNV hot spot areas, our study affirmed the importance of birds and horses as sentinels for human WNV-infections. Thus, information dissemination regarding WNV-infections should be conducted among healthcare and veterinary workers and the greater public. Furthermore, we recommend that horses located in these WNV hotspot areas and nearby regions be vaccinated against WNV according to the recommendations of the Standing Committee on Vaccination for Veterinary Medicine in Germany (StIKo Vet).

## Acknowledgements

We are grateful to Patrick Zitzow, Cornelia Steffen, Katja Wittig, and Katrin Schwabe for excellent technical assistance. We are indebted to Dr. Susanne Fischer for the significant input regarding affinity propagation clustering. We thank Sabine Bock (Berlin-Brandenburg State Laboratory, Frankfurt (Oder)), Kerstin Albrecht (State Institute for Consumer Protection of Saxony-Anhalt, Department of Veterinary Medicine, Stendal), Andrea Konrath (Saxon State Laboratory of Health and Veterinary Affairs, Leipzig), Aemero Muluneh (Saxon State Laboratory of Health and Veterinary Affairs, Dresden), Timo Siempelkamp (Thuringia Office for Consumer Protection, Bad Langensalza), Michael Sieg (Institute of Virology, Faculty of Veterinary Medicine, Leipzig University, Leipzig), Claudia Szentiks (Leibniz Institute for Zoo and Wildlife Research, Berlin), Dominik Fischer (Clinic for Birds, Reptiles, Amphibians and Fish, Justus Liebig University Giessen, Giessen), Claudia Sauerwald (Landesbetrieb Hessisches Landeslabor, Veterinary Virology and Molecular Biology, Gießen) and the other colleagues of the veterinary authorities and veterinary laboratories of the federal states for the supply of the samples and we are very grateful for the continuous support. Furthermore, we thank the staff of the different bird clinics, rehabilitation centers, zoological gardens and “Tierparks” of Germany as partners in the nation-wide wild bird surveillance network for zoonotic arthropod-borne viruses, which collected and sent samples for the present study. Furthermore, we want to thank Patrick Wysocki and Daike Lehnau (Institute of Epidemiology, FLI, Greifswald-Insel Riems) for their help by producing the epidemiological datasets and Jacqueline King (Institute of Diagnostic Virology, FLI, Greifswald-Insel Riems) for proofreading the manuscript.

This work was funded by European Union’s Horizon 2020 research and innovation program under the Marie Skłodowska-Curie Actions grant agreement no. 721367 (HONOURs) and in part by the EU Horizon 2020 program grant agreement no. 874735 (VEO) and by the German Federal Ministry of Food and Agriculture (BMEL) through the Federal Office for Agriculture and Food (BLE), grant number 2819113919 (CuliFo2) as well as by the German Center for Infection Research (DZIF) under project number TTU 01.804 (WBA-Zoo), and by the Federal Ministry of Education and Research within the research consortium ‘‘ZooBoCo” (Grant No. 01KI1722A).

## Data availability

The nucleotide sequences from this study are available from the INSDC databases study accession PRJEB47687.

## Supplementary material

### R-Code

\### AP-Clustering of genome sequences based on an identity-matrix

\## load necessary R-packages library(apcluster) library(writexl)

\## prepare workspace

rm(list = ls()) # clean up workspace to prevent from interference between calculation and pre- existing data

\## settings; when multiple values are provided, all possible combinations are tested

minStepNumbers <- c(1000, 2000, 5000, 10000) # the minimum number of steps to divide the input preference range for plateau calculations

stepFactor <- 10 # currently not used; Factor to calculate th allowed maximum step number from the used minStepNumbers

windowProportions <- c(0.01, 0.005, 0.0025, 0.001, 0.0001) # the fraction of the complete steps (see above) the partial dataset used for the APC should have

minPlateauWindows <- 3 # the allowed minimum the actual window may have (values < 3 do not make sense)

groupSizes <- c(5, 7, 10) # the minimum number of sequences of a subgroup to use as input for a further subgrouping

maxGroupDepth <- 5 # number of hierarchy levels to calculate with 1=Lineage/2=Clade/3=Subclade/4=Cluster/5=Subcluster

\## load and prepare input data

setwd(“M:/R-work/PS_wnv_apc/testWL2/”) # choose the folder to read the input from and to save the output

dateipfad <- “M:/R-work/PS_wnv_apc/testWL2/WL2_R01_cleaned.csv” # choose the input file

sequenceIdentityMatrix <- read.csv(dateipfad, row.names = 1) # read in the matrix of sequence identities

if(identical(colnames(sequenceIdentityMatrix), rownames(sequenceIdentityMatrix))) { # check whether or not the dimnames are identical, if not do not calculate because it might generate invalid results

dissimilarityMatrix <- -1*((1-as.matrix(sequenceIdentityMatrix))^2) # convert the sequence identity matrix into a matrix of dissimilarities as this is the input for affinity propagation clustering

for(aktStepNumber in minStepNumbers) { # iterate the calculations over all preset numbers of steps over the input preference range

for(winProp in windowProportions) { # iterate the calculations over all preset proportions for plateau definition

for(aktGroupSize in groupSizes) { # iterate the calculations over all preset minimum group sizes

for(aktPlateauWindow in minPlateauWindows) {

dateien <- list.files() # read in a list of files currently present in your target folder ## prepare necessary objects to accomodate results

paramCollection <- data.frame(level = NA, group = NA, inputGroupSize = NA, usedStepWidth = NA, setIterations = NA, terminatingInPref = NA,

usedIterations = NA, minPrefRange = NA, maxPrefRange = NA,

defaultStepNumber = NA, lastStable = NA, longest = NA, defaultAPC = NA, cutCrit = NA, usedWindowSize = NA) # set up a data.frame to collect all

relevant parameters used in the current iteration

allResults <- vector(“list”, 4) # set up a list to collect all clustering results from the current iteration

names(allResults) <- c(“group”, “defaultAPresult”, “cutreeResult”, “aggExResult”) # name the elements of the list

\## calculate settings adjusted according to above input

maxStepNumber <- aktStepNumber * stepFactor # currently not used; calculate the maximum allowed step number

initSteps <- (aktStepNumber + maxStepNumber) / 2 # currently not used; calculate the initial number of steps to start with

plateauWindow <- floor(aktStepNumber * winProp) # set the window size to fit the actual step number and interrogate always the same portion of the overall range

if(plateauWindow < aktPlateauWindow) plateauWindow <- aktPlateauWindow ## in case the combined settings result in a window smaller than the minimum allowed window size, adjust the setting to the allowed minimum

\## prepare filenames to save results

datensatz <- sub(“\\.[[:alpha:]]{1,}$”, “”, basename(dateipfad)) # extract the name of the dataset from the filepath

filename <- paste(datensatz, “.test.Affiliations-minSteps_”, aktStepNumber, “-window_”, plateauWindow,

”-minMembers_”, aktGroupSize, “-maxGroupDepth_”, maxGroupDepth, “.xlsx”, sep

= “”) # construct filename containing the distinguishing parameters

if(is.element(filename, dateien) == FALSE) { # check whether the calculation using the recent parameter combination was already initiated; only if not, continue calculating, otherwise skip to the next combination

write_xlsx(data.frame(“Analysis in progress”), filename, col_names = FALSE) # write a file into the current folder to mark the parameter combination in progress

print(paste(“Analysis in progress: “, filename, sep = “”)) # output user information

seqSubset <- colnames(dissimilarityMatrix) # define the initial set of sequences, i.e. use all sequences of the dataset

affiliations <- data.frame(matrix(nrow = length(seqSubset), ncol = maxGroupDepth + 3)) # set up an object to save the results of the APC; needs 3 more columns than the number of hierarchy levels defined by maxGroupDepth

colnames(affiliations) <- c(“accession”, “affiliation”, paste(“affil”, 0:maxGroupDepth, sep = “0”)) # set the column names

rownames(affiliations) <- colnames(dissimilarityMatrix) # set rownames identical with colnames of the input identity matrix

affiliations$accession <- rownames(affiliations) # copy the rownames to the first column of the result matrix

affiliations$affil00 <- 1 # set the initial affiliation level to 1 for all sequences affiliations$affiliation <- 0 # set the affiliation 0 for all sequences

for(mgd in 0:(maxGroupDepth - 1)) { # iteratively run through the grouping for all sequences for the given number of hierarchy levels

prevLevel <- paste(“affil”, mgd, sep = “0”) # the previous hierarchy level, the starting point for subsetting the dataset

currLevel <- paste(“affil”, (mgd + 1), sep = “0”) # the current hierarchy level to be determined for the respective subset of sequences

affiliations[, currLevel] <- 0 # set the initial affiliations in the currently analysed hierarchy level to the default value

for(aktSubGroup in unique(affiliations$affiliation[affiliations[,prevLevel] > 0])) { # run the grouping for the current hierarchy level for all subgroups of the preceding level

seqSubset <- affiliations$accession[affiliations$affiliation == aktSubGroup] # generate the list of sequence names belonging to the currently analysed subgroup

if(length(seqSubset) < aktGroupSize) affiliations[seqSubset, currLevel] <- -1 else { # check whether the number of sequences in the current group is sufficient according to the preset minimum group size to allow for further subdivision; if not, set the current affiliation -1 to stop further evaluation in the subsequent iterations

workmat <- as.matrix(dissimilarityMatrix[seqSubset, seqSubset]) # if the current group size allows for further subdivision, get the working matrix only containing data of the relevant subset

workmatAPC <- apcluster(workmat, details=TRUE, q=0.5) # calculate the AP-clustering of the current data subset using the default input preference q

prefRang <- preferenceRange(workmat, exact=TRUE) # determine the input preference range of the data subset

pStepWidth <- abs((prefRang[2] - prefRang[1]) / (aktStepNumber - 2)) # adjust the step width to cover the complete input preference range in equal steps

inPref <- unique(c(prefRang[1], seq(prefRang[1], prefRang[2], pStepWidth), prefRang[2])) # calculate all input preferences to use in the APC iterations

plateauWindow <- floor(length(inPref) * winProp) # set the window size to fit the actual step number and interrogate always the same portion of the overall range

if(plateauWindow < aktPlateauWindow) plateauWindow <- aktPlateauWindow # in case the calculated window size for the determination of a cluster number plateau is lower than the preset lower level, adjust the size of the window used to define the plateau to fit with the lower limit

if(length(inPref) > plateauWindow) { # test whether sufficient iterations are performed to cover the set window for plateau determination, only start iterating if yes because otherwise an error will occur

clusTab <- data.frame(inPref, numClust = NA, windowStDev = NA, windowMean = NA, increase = TRUE, stDevOK = TRUE) # prepare table to save the results of all clustering iterations to enable testing whether or not the stopping criteria are met

i <- 0 # define counter

stopAPC <- FALSE # set the control variable

while(i < nrow(clusTab) & stopAPC == FALSE) { # repeat the calculations for AP clustering of the current data subset until either of the stopping criteria is met

i <- i + 1 # increase counter for current iteration

j<-apcluster(workmat, p = clusTab$inPref[i]) # determine the number of AP clusters in the data subset with the given input preference (as previously defined from the preference range and chosen number of iterations)

clusTab$numClust[i] <- length(j@clusters) # record the number of AP clusters corresponding with the input preference

if(i <= plateauWindow) { # check whether or not sufficient data for calculation of mean and SD from number of clusters is available

if(i == 1) clusTab$windowStDev[i] <- 0 else clusTab$windowStDev[i] <- sd(clusTab$numClust[1:i]) # if not, set SD of cluster number within window 0 in case of first iteration, otherwise adjust SD calculation to available data instead of preset window

clusTab$windowMean[i] <- mean(clusTab$numClust[1:i]) # calculate mean from the available data definition

} else { # number of performed iterations higher than window size for plateau clusTab$windowStDev[i] <- sd(clusTab$numClust[(i - plateauWindow + 1):i]) #

calculate SD of cluster number from recent and preceding iterations (in total preset number of

iterations)

clusTab$windowMean[i] <- mean(clusTab$numClust[(i - plateauWindow + 1):i]) # calculate mean cluster number from recent and preceding iterations (in total preset number of iterations)

clusTab$increase[i] <- clusTab$windowMean[i] >= clusTab$windowMean[(i-1)] # test whether the number of clusters is the same as or larger than in the preceding iteration, because a decrease is deemed a disruption and leads to termination of the iterative AP clustering

tempTab <- clusTab[(i - plateauWindow + 1):i,] # make subset of the table only containing data of the plateauWindow number of rows including the last iteration

clusTab$stDevOK[i] <- nrow(tempTab[tempTab$windowStDev != 0,]) < plateauWindow # test whether the SD of the cluster number returns to 0 after an increase of the cluster number (this must be the case if the cluster number is stable for at least plateauWindow iterations), if not a disruption occurred

stopAPC <- !(clusTab$increase[i] == TRUE & clusTab$stDevOK[i] == TRUE) # check whether or not both criteria to enter the next iteration, i.e. not to terminate the loop, are fulfilled

}

}

setIterations <- nrow(clusTab) # record the set maximum number of iterations

clusTab <- clusTab[1:(i - plateauWindow),] # cut the table to the number of iterations before the disruption occurred

usedIterations <- nrow(clusTab) # record the number of iterations with stable plateau lastStable <- clusTab$numClust[nrow(clusTab)] # record the number clusters in the last stable plateau

firstPlateau <- min(clusTab$numClust) # record the number of clusters in the first observed plateau

plateauSummary <- clusTab$numClust # prepare the identification of the longest plateau

plateauSummary <- plateauSummary[plateauSummary > firstPlateau] # only use values higher than the first plateau (as per the definition in Susanne Fischer’s paper the first plateau is not valid)

plateauSummary <- summary(as.factor(plateauSummary), maxsum = length(unique(plateauSummary))) # summarize how often each number of clusters was observed to define the longest plateau, i.e. the number of clusters that was most often observed before the disruption

if(is.element(TRUE, duplicated(plateauSummary))) longestPlateau <- as.numeric(names(plateauSummary[plateauSummary == max(plateauSummary)])) else longestPlateau <- as.numeric(names(which.max(plateauSummary))) # determine the number of clusters constituting the longest plateau; in case 2 or more cluster numbers are present the same number of iterations, use the higher number of clusters in order not to reduce the cluster number too stringently

allPlateaus <- c(lastStable, longestPlateau) # concatenate the determined clusternumbers from the longest and the last stable plateau

if(is.element(TRUE, allPlateaus < length(workmatAPC@clusters))) { # define the best number of clusters to use and record the used choice; the best choice is the highest number of clusters that is equal or lower than the number of clusters determined with the default input preference, therefore, test whether either the last stable or the longest plateau are more stringent than the default

cutNum <- max(allPlateaus[allPlateaus < length(workmatAPC@clusters)]) # record the number of clusters to use for cutting the tree (below)

if(cutNum == lastStable) cutCrit <- “last” else cutCrit <- “longest” # record the choice in case the default is replaced

} else { # the default value is used

cutNum <- length(workmatAPC@clusters) # record the default value of the cluster number to use it for tree cutting below

cutCrit <- “defaultAPC” # record the used choice

}

aggdissimilarityMatrix <- aggExCluster(workmat, workmatAPC) # agglomerative hierarchical clustering

grouping <- cutree(aggdissimilarityMatrix, k = cutNum) # cutting the tree to determine the resulting grouping of sequences; k = number of groups to have = cluster number as determined above

if(length(grouping@clusters) == 1) for(g in 1:length(grouping@clusters)) affiliations[rownames(workmat)[grouping@clusters[[g]]], currLevel] <- -3 else for(g in 1:length(grouping@clusters)) affiliations[rownames(workmat)[grouping@clusters[[g]]], currLevel] <- g # record the grouping in the current subset of the current hierarchical level; in case the subset cannot be further subdivided (number of clusters is 1), record -3 to label the grouping being terminated for the subset because it cannot be further subdivided; in all other cases, record the group affiliations per sequence

} else affiliations[seqSubset, currLevel] <- -2 # in case there is not enough steps for the calculations, report -2 to label the subgroup for subsequent cycles and error analysis

}

\## in the following lines, record all current settings of the iteration paramCollection$terminatingInPref[nrow(paramCollection)] <-

clusTab$inPref[nrow(clusTab)]

paramCollection$level[nrow(paramCollection)] <- mgd

paramCollection$group[nrow(paramCollection)] <- aktSubGroup paramCollection$inputGroupSize[nrow(paramCollection)] <- length(seqSubset) paramCollection$usedStepWidth[nrow(paramCollection)] <- pStepWidth paramCollection$setIterations[nrow(paramCollection)] <- setIterations paramCollection$usedIterations[nrow(paramCollection)] <- usedIterations paramCollection$minPrefRange[nrow(paramCollection)] <- prefRang[1] paramCollection$maxPrefRange[nrow(paramCollection)] <- prefRang[2] paramCollection$lastStable[nrow(paramCollection)] <- lastStable

if(length(longestPlateau > 0)) paramCollection$longest[nrow(paramCollection)] <- longestPlateau else paramCollection$longest[nrow(paramCollection)] <- NA

paramCollection$defaultAPC[nrow(paramCollection)] <- length(workmatAPC@clusters) paramCollection$cutCrit[nrow(paramCollection)] <- cutCrit paramCollection$usedWindowSize[nrow(paramCollection)] <- plateauWindow

paramCollection <- rbind(paramCollection, NA) # add the next line to the table to accommodate the data of the next iteration

\## End parameter recording

\## in the following lines, record all results of the current iteration allResults$group <- append(allResults$group, aktSubGroup) allResults$defaultAPresult <- append(allResults$defaultAPresult, workmatAPC) allResults$aggExResult <- append(allResults$aggExResult, aggdissimilarityMatrix) allResults$cutreeResult <- append(allResults$cutreeResult, grouping)

\## End results recording

}

if(mgd == 0) affiliations$affiliation[affiliations[, currLevel] > 0] <- affiliations[affiliations[, currLevel] > 0, currLevel] else affiliations$affiliation[affiliations[, currLevel] > 0] <- paste(affiliations$affiliation[affiliations[, currLevel] > 0], affiliations[affiliations[, currLevel] > 0, currLevel], sep = “.”) # construct the overall group designation from the previously present portion and the currently analyzed hierarchy level; in case it is the first level iteration, replace the present values with the current

}

affiliations$affil00 <- NULL # delete the initial grouping

\## save results to disk write_xlsx(affiliations, filename)

write_xlsx(paramCollection, sub(“Affiliations”, “usedParameters”, filename)) save.image(file = sub(“Affiliations”, “CompleteData”, sub(“xlsx”, “RData”, filename)))

}

}

}

}

}

} else print(“Please check the column and row names in your input file! They must be identical!”)

---

**Figure S1.**
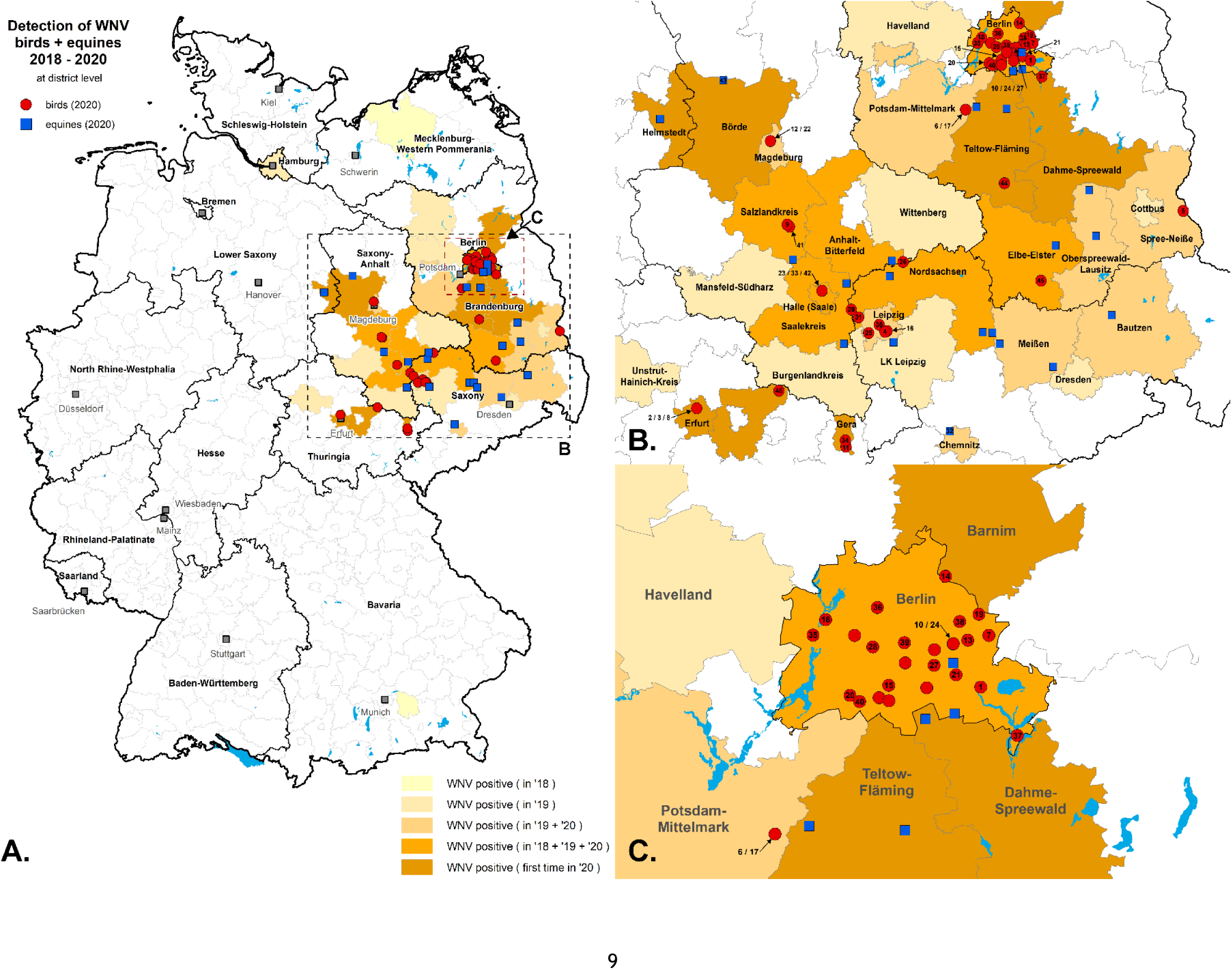
Geographic distribution of WNV cases in Germany in 2020 (depicted on district level) as shown in A. Specific areas with WNV cases in the areas of Saxony, Saxony-Anhalt, Thuringia, Berlin and Brandenburg were shown in B and WNV cases in Berlin and surrounding areas in Brandenburg were shown in C. Blue squares and red circles indicate notifiable WNV cases of horses and birds. WNV cases with numbers indicated that these samples were subjected to whole-genome sequencing. WNV cases that were not selected for sequencing (e.g., IgM-positive cases or high C_q_ values) remain unnumbered. Intensity of the colored background at district level indicates the frequency, how often an area was affected by WNV activity in prior years.

**Figure S2.**
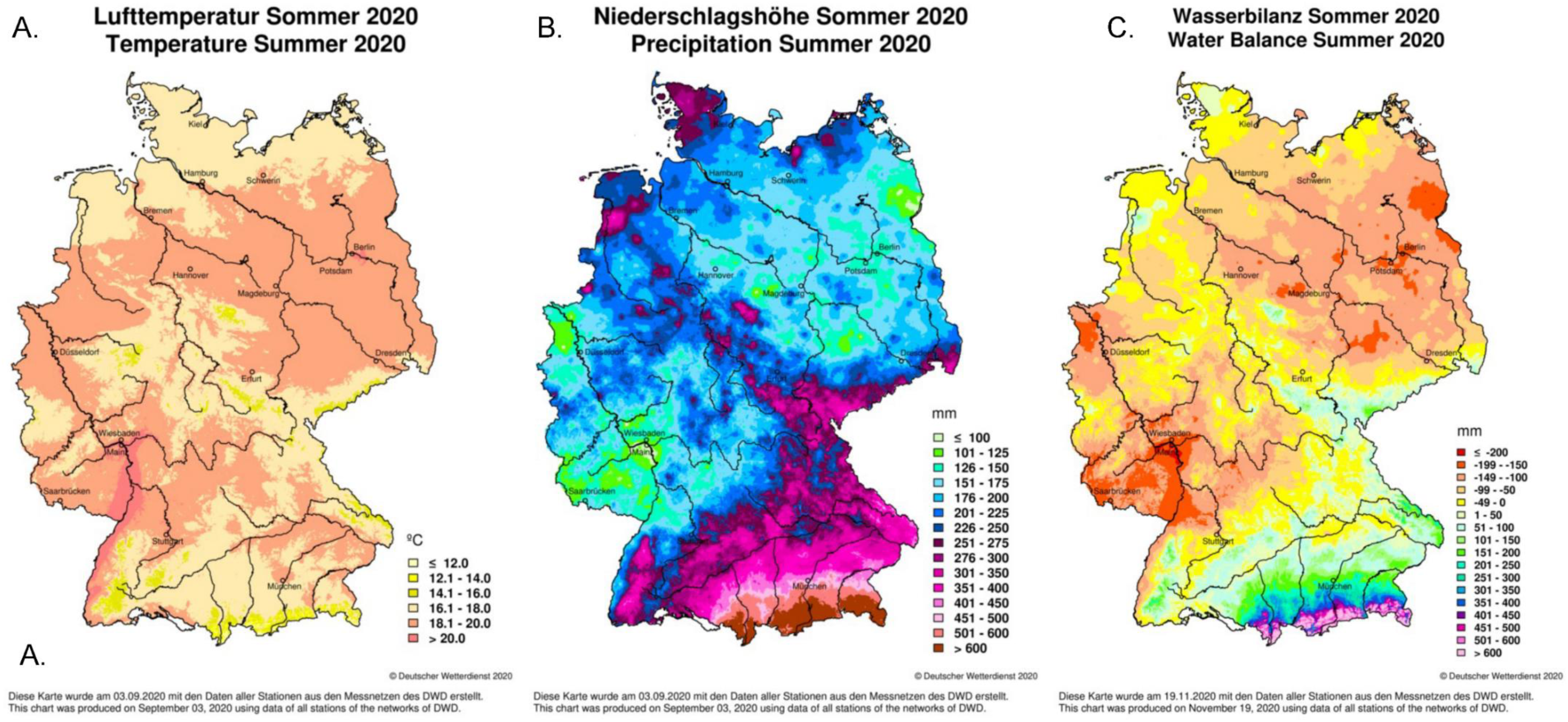
Climatological maps of Germany displaying A) temperature (in degree celsius), B) precipitation (in millimeter) and C) water balance (in millimeter) based on data collected in summer 2020. Climatological maps were downloaded from Deutscher Wetterdienst; https://www.dwd.de/EN/climate_environment/climatemonitoring/germany/germany_node.html (Deutscher Wetterdienst 2020)

**Table S1.**
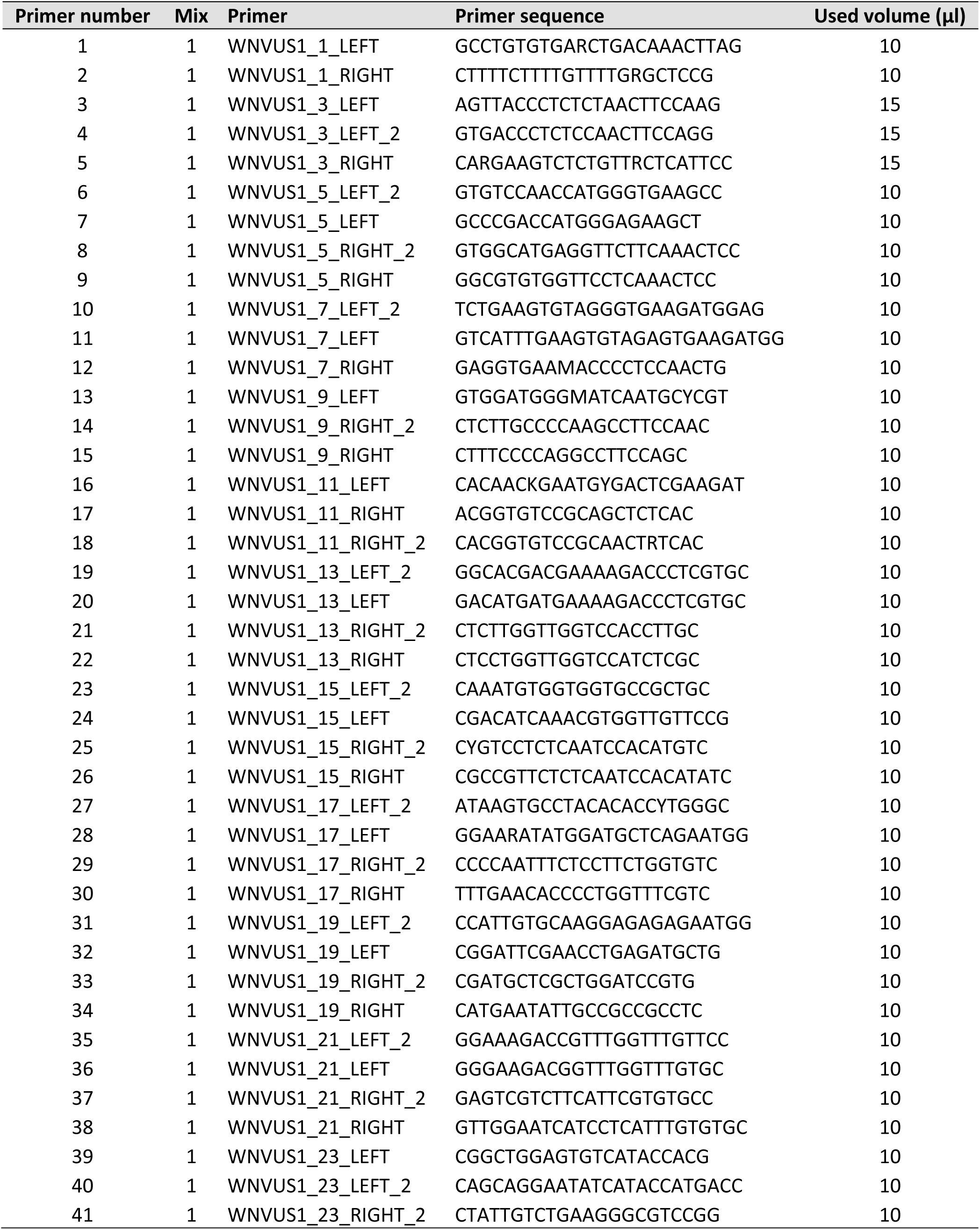
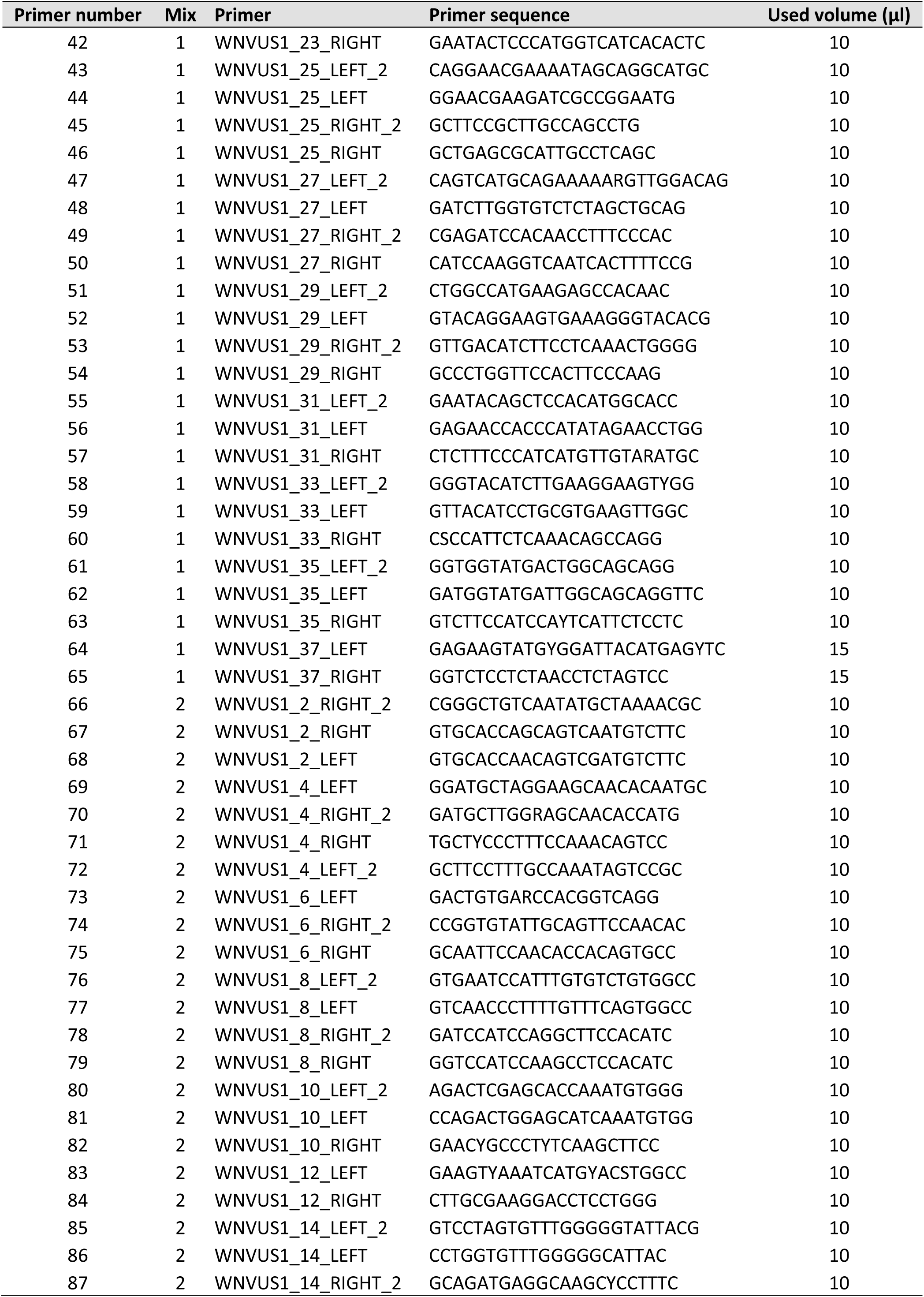
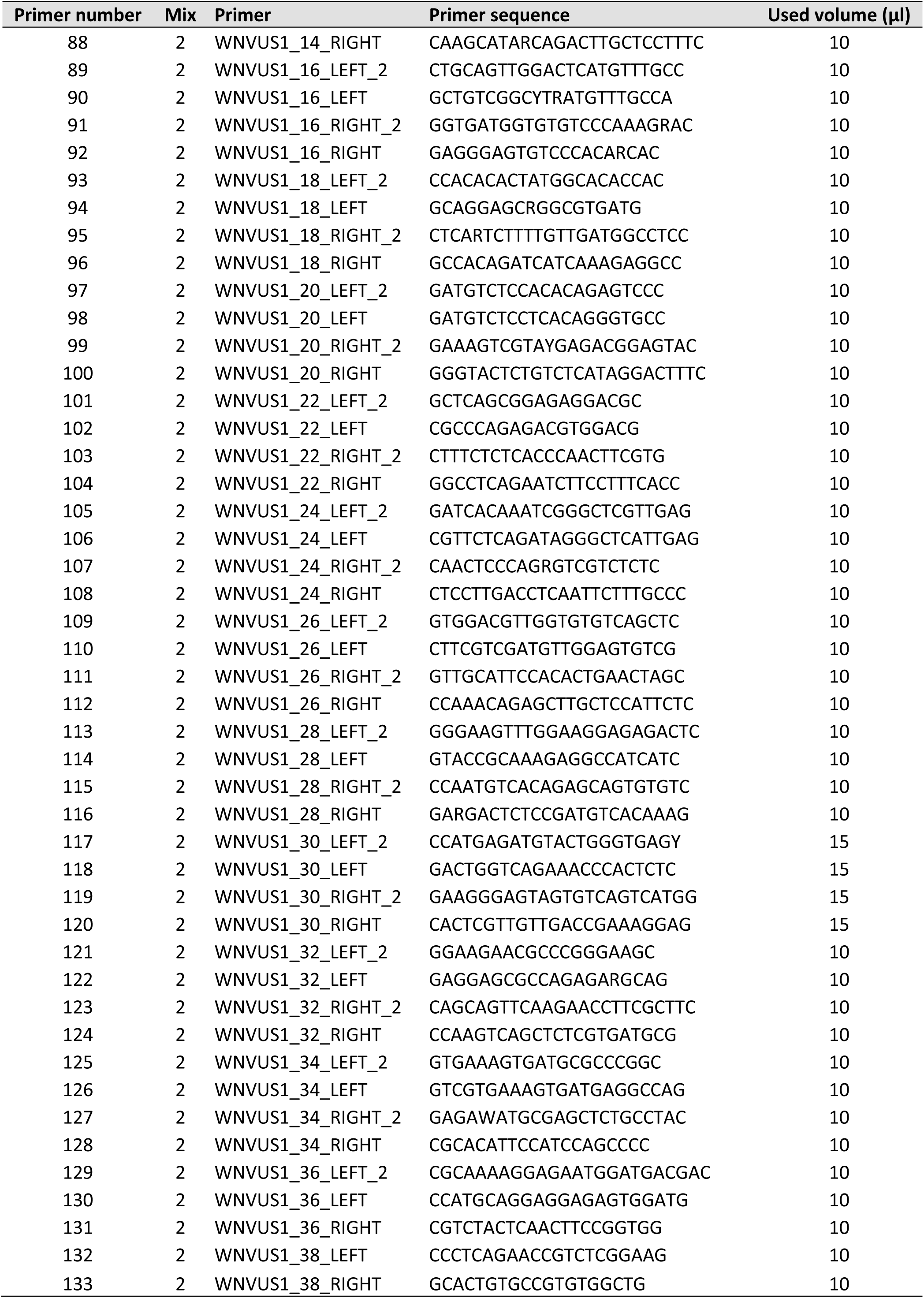
West Nile Virus primer sequences from Sikkema et al. 2020. Each primer stock was normalized to 100 micromolar concentration. Volumes of each primer per primer mix (mix 1 or 2) were specified in the table below. These primers mixes were then subjected to 1:10 dilution.

**Table S2.** Samples used to validate the WNV multiplex PCR High-throughput sequencing (HTS) in comparison with the result of unbiased and direct HTS approach.

| Sample |  |  | Reference sequence |  |  |  | This study |  |  | Result of<br>sequence<br>comparison |
| --- | --- | --- | --- | --- | --- | --- | --- | --- | --- | --- |
| Code | ID | Cq value | Library ID | applied protocol | INSDC<br>Accesssion | Length (nt) | Library ID | INSDC<br>Accesssion | Length (nt) |  |
| C1 | ED-I-62/19 | 22.9 | lib03378 | Wylezich et al 2018 | LR743425 | 11,060 | lib04562 | XX | 10,989 | identical |
| C2 | ED-I-156/19 | 11.8 | lib03418 | Wylezich et al 2018 | LR743423 | 11,027 | lib04563 | XX | 10,989 | Identical |
| C3 | ED-I-155/19 | 17.5 | lib03420 | Wylezich et al 2018 | LR743422 | 11,010 | lib04564 | XX | 10,987 | Identical |
| C4 | ED-I-115/19 | 31.5 | lib03988 | Wylezich et al 2021 | LR989891 | 6,470 | lib04565 | XX | 10,989 | 8 substitutions<sup>1</sup> |
| C5 | ED-I-127-18 | 29.4 | lib03224 | Wylezich et al 2021 | Unpublished | 6,786 | lib04748 | XX | 10,988 | 2 substitutions<sup>1</sup> |

<sup>1</sup> based on available partial reference sequence; insertions and deletions not considered

**Table S3.** List of full genome sequences retrieved from Genbank

| Accession number | Used in Dataset |
| --- | --- |
| DQ116961 | WL2 |
| HQ537483 | WL2 |
| JN858070 | WL2 |
| KC407673 | WL2 |
| KC496015 | WL2 |
| KC496016 | WL2 |
| KF179639 | WL2 |
| KF179640 | WL2 |
| KF588365 | WL2 |
| KF647249 | WL2 |
| KF647250 | WL2 |
| KF647251 | WL2 |
| KF647252 | WL2 |
| KF823806 | WL2 |
| KJ577738 | WL2 |
| KJ577739 | WL2 |
| KJ883342 | WL2 |
| KJ883343 | WL2 |
| KJ883344 | WL2 |
| KJ883345 | WL2 |
| KJ883346 | WL2 |
| KJ883348 | WL2 |
| KJ883349 | WL2 |
| KJ883350 | WL2 |
| KM203860 | WL2 |
| KM203861 | WL2 |
| KM203862 | WL2 |
| KM203863 | WL2 |
| KM659876 | WL2 |
| KP109691 | WL2 |
| KP109692 | WL2 |
| KP780837 | WL2 |
| KP780838 | WL2 |
| KP780839 | WL2 |
| KP789953 | WL2 |
| KP789954 | WL2 |
| KP789955 | WL2 |
| KP789956 | WL2 |
| KP789957 | WL2 |
| KP789958 | WL2 |
| KP789959 | WL2 |
| KP789960 | WL2 |
| KT207792 | WL2 |
| KT359349 | WL2 |
| KT757318 | WL2 |
| KT757319 | WL2 |
| KT757320 | WL2 |
| KT757321 | WL2 |
| KT757322 | WL2 |
| KT757323 | WL2 |
| KU206781 | WL2 |
| KU573080 | WL2 |
| KU573081 | WL2 |
| KU573082 | WL2 |
| KU573083 | WL2 |
| KX375812 | WL2 |
| KY594040 | WL2 |
| LR743421 | WL2 |
| LR743422 | WL2 |
| LR743423 | WL2 |
| LR743424 | WL2 |
| LR743425 | WL2 |
| LR743426 | WL2 |
| LR743427 | WL2 |
| LR743428 | WL2 |
| LR743429 | WL2 |
| LR743430 | WL2 |
| LR743431 | WL2 |
| LR743432 | WL2 |
| LR743433 | WL2 |
| LR743434 | WL2 |
| LR743435 | WL2 |
| LR743436 | WL2 |
| LR743437 | WL2 |
| LR743442 | WL2 |
| LR743443 | WL2 |
| LR743444 | WL2 |
| LR743445 | WL2 |
| LR743446 | WL2 |
| LR743447 | WL2 |
| LR743448 | WL2 |
| LR743449 | WL2 |
| LR743450 | WL2 |
| LR743451 | WL2 |
| LR743452 | WL2 |
| LR743453 | WL2 |
| LR743454 | WL2 |
| LR743455 | WL2 |
| LR743456 | WL2 |
| LR743457 | WL2 |
| LR743458 | WL2 |
| LR989885 | WL2 |
| LR989888 | WL2 |
| MF984337 | WL2 |
| MF984338 | WL2 |
| MF984339 | WL2 |
| MF984340 | WL2 |
| MF984341 | WL2 |
| MF984342 | WL2 |
| MF984343 | WL2 |
| MF984344 | WL2 |
| MF984345 | WL2 |
| MF984346 | WL2 |
| MF984347 | WL2 |
| MF984348 | WL2 |
| MF984349 | WL2 |
| MF984350 | WL2 |
| MF984351 | WL2 |
| MF984352 | WL2 |
| MH021189 | WL2 |
| MH244510 | WL2 |
| MH244511 | WL2 |
| MH244512 | WL2 |
| MH244513 | WL2 |
| MH549209 | WL2 |
| MH910045 | WL2 |
| MH924836 | WL2 |
| MH986055 | WL2 |
| MH986056 | WL2 |
| MK473443 | WL2 |
| MK947396 | WL2 |
| MK947397 | WL2 |
| MN480792 | WL2 |
| MN480793 | WL2 |
| MN480794 | WL2 |
| MN480795 | WL2 |
| MN481589 | WL2 |
| MN481590 | WL2 |
| MN481591 | WL2 |
| MN481592 | WL2 |
| MN481593 | WL2 |
| MN481594 | WL2 |
| MN481595 | WL2 |
| MN481596 | WL2 |
| MN481597 | WL2 |
| MN652878 | WL2 |
| MN652879 | WL2 |
| MN652880 | WL2 |
| MN794935 | WL2 |
| MN794937 | WL2 |
| MN794938 | WL2 |
| MN794939 | WL2 |
| MN939557 | WL2 |
| MN939558 | WL2 |
| MN939559 | WL2 |
| MN939560 | WL2 |
| MN939561 | WL2 |
| MN939562 | WL2 |
| MN939562 | WL2 |
| MN939564 | WL2 |
| MT341470 | WL2 |
| MT341471 | WL2 |
| MT341472 | WL2 |
| MT863560 | WL2 |
| MT863561 | WL2 |
| MW036634 | WL2 |
| MW142223 | WL2 |
| MW142224 | WL2 |
| MW142226 | WL2 |
| MW142227 | WL2 |
| AF196835 | TD01 / TD03 |
| AY277251 | TD01 / TD03 |
| AY532665 | TD01 / TD03 |
| AY701412 | TD01 / TD03 |
| AY701413 | TD01 / TD03 |
| AY765264 | TD01 / TD03 |
| DQ176636 | TD01 / TD03 |
| DQ256376 | TD01 / TD03 |
| DQ786573 | TD01 / TD03 |
| EF429197 | TD01 / TD03 |
| EF429198 | TD01 / TD03 |
| EF429199 | TD01 / TD03 |
| EF429200 | TD01 / TD03 |
| EU082200 | TD01 / TD03 |
| EU249803 | TD01 / TD03 |
| FJ159129 | TD01 / TD03 |
| FJ159130 | TD01 / TD03 |
| FJ159131 | TD01 / TD03 |
| FJ425721 | TD01 / TD03 |
| FJ483548 | TD01 / TD03 |
| FJ483549 | TD01 / TD03 |
| FJ766331 | TD01 / TD03 |
| FJ766332 | TD01 / TD03 |
| GQ379161 | TD01 / TD03 |
| GQ851602 | TD01 / TD03 |
| GQ851603 | TD01 / TD03 |
| GQ851604 | TD01 / TD03 |
| GQ851605 | TD01 / TD03 |
| GQ851606 | TD01 / TD03 |
| GQ851607 | TD01 / TD03 |
| GQ903680 | TD01 / TD03 |
| GU011992 | TD01 / TD03 |
| HM051416 | TD01 / TD03 |
| HM147822 | TD01 / TD03 |
| HM147823 | TD01 / TD03 |
| HM147824 | TD01 / TD03 |
| HM152775 | TD01 / TD03 |
| HQ537483 | TD01 / TD03 |
| JF707789 | TD01 / TD03 |
| JF719066 | TD01 / TD03 |
| JF719067 | TD01 / TD03 |
| JF719068 | TD01 / TD03 |
| JF719069 | TD01 / TD03 |
| JN393308 | TD01 / TD03 |
| JN858069 | TD01 / TD03 |
| JN858070 | TD01 / TD03 |
| JQ928174 | TD01 / TD03 |
| JQ928175 | TD01 / TD03 |
| JX041628 | TD01 / TD03 |
| JX041629 | TD01 / TD03 |
| JX041630 | TD01 / TD03 |
| JX041632 | TD01 / TD03 |
| JX041634 | TD01 / TD03 |
| JX123030 | TD01 / TD03 |
| JX123031 | TD01 / TD03 |
| JX442279 | TD01 / TD03 |
| JX556213 | TD01 / TD03 |
| KC407673 | TD01 / TD03 |
| KC496015 | TD01 / TD03 |
| KC496016 | TD01 / TD03 |
| KC601756 | TD01 / TD03 |
| KC954092 | TD01 / TD03 |
| KF179639 | TD01 / TD03 |
| KF179640 | TD01 / TD03 |
| KF234080 | TD01 / TD03 |
| KF647251 | TD01 / TD03 |
| KF647253 | TD01 / TD03 |
| KJ831223 | TD01 / TD03 |
| KJ883346 | TD01 / TD03 |
| KJ934710 | TD01 / TD03 |
| KM052152 | TD01 / TD03 |
| KM203861 | TD01 / TD03 |
| KM203862 | TD01 / TD03 |
| KM203863 | TD01 / TD03 |
| KP109692 | TD01 / TD03 |
| KP780837 | TD01 / TD03 |
| KP780838 | TD01 / TD03 |
| KP780839 | TD01 / TD03 |
| KP780840 | TD01 / TD03 |
| KT163243 | TD01 / TD03 |
| KT207791 | TD01 / TD03 |
| KT207792 | TD01 / TD03 |
| KT359349 | TD01 / TD03 |
| KT934796 | TD01 / TD03 |
| KT934797 | TD01 / TD03 |
| KT934798 | TD01 / TD03 |
| KT934799 | TD01 / TD03 |
| KT934800 | TD01 / TD03 |
| KT934801 | TD01 / TD03 |
| KT934802 | TD01 / TD03 |
| KT934803 | TD01 / TD03 |
| KU588135 | TD01 / TD03 |
| KY703854 | TD01 / TD03 |
| KY703855 | TD01 / TD03 |
| KY703856 | TD01 / TD03 |
| AF196835 | TD02 / TD03 |
| AF202541 | TD02 / TD03 |
| AF206518 | TD02 / TD03 |
| AF260967 | TD02 / TD03 |
| AF260968 | TD02 / TD03 |
| AF260969 | TD02 / TD03 |
| AF317203 | TD02 / TD03 |
| AF404753 | TD02 / TD03 |
| AF404754 | TD02 / TD03 |
| AF404755 | TD02 / TD03 |
| AF404756 | TD02 / TD03 |
| AF404757 | TD02 / TD03 |
| AF481864 | TD02 / TD03 |
| AF533540 | TD02 / TD03 |
| AJ965626 | TD02 / TD03 |
| AJ965628 | TD02 / TD03 |
| AM404308 | TD02 / TD03 |
| AY262283 | TD02 / TD03 |
| AY268132 | TD02 / TD03 |
| AY268133 | TD02 / TD03 |
| AY274504 | TD02 / TD03 |
| AY277252 | TD02 / TD03 |
| AY278441 | TD02 / TD03 |
| AY278442 | TD02 / TD03 |
| AY289214 | TD02 / TD03 |
| AY603654 | TD02 / TD03 |
| AY646354 | TD02 / TD03 |
| AY660002 | TD02 / TD03 |
| AY701412 | TD02 / TD03 |
| AY701413 | TD02 / TD03 |
| AY712945 | TD02 / TD03 |
| AY712946 | TD02 / TD03 |
| AY712947 | TD02 / TD03 |
| AY712948 | TD02 / TD03 |
| AY795965 | TD02 / TD03 |
| DQ005530 | TD02 / TD03 |
| DQ080051 | TD02 / TD03 |
| DQ080052 | TD02 / TD03 |
| DQ080053 | TD02 / TD03 |
| DQ080054 | TD02 / TD03 |
| DQ080055 | TD02 / TD03 |
| DQ080056 | TD02 / TD03 |
| DQ080057 | TD02 / TD03 |
| DQ080058 | TD02 / TD03 |
| DQ080059 | TD02 / TD03 |
| DQ080060 | TD02 / TD03 |
| DQ080061 | TD02 / TD03 |
| DQ080062 | TD02 / TD03 |
| DQ080063 | TD02 / TD03 |
| DQ080064 | TD02 / TD03 |
| DQ080065 | TD02 / TD03 |
| DQ080066 | TD02 / TD03 |
| DQ080067 | TD02 / TD03 |
| DQ080068 | TD02 / TD03 |
| DQ080069 | TD02 / TD03 |
| DQ080070 | TD02 / TD03 |
| DQ080071 | TD02 / TD03 |
| DQ080072 | TD02 / TD03 |
| DQ118127 | TD02 / TD03 |
| DQ164186 | TD02 / TD03 |
| DQ164187 | TD02 / TD03 |
| DQ164188 | TD02 / TD03 |
| DQ164189 | TD02 / TD03 |
| DQ164190 | TD02 / TD03 |
| DQ164191 | TD02 / TD03 |
| DQ164192 | TD02 / TD03 |
| DQ164193 | TD02 / TD03 |
| DQ164194 | TD02 / TD03 |
| DQ164195 | TD02 / TD03 |
| DQ164196 | TD02 / TD03 |
| DQ164197 | TD02 / TD03 |
| DQ164198 | TD02 / TD03 |
| DQ164199 | TD02 / TD03 |
| DQ164200 | TD02 / TD03 |
| DQ164201 | TD02 / TD03 |
| DQ164202 | TD02 / TD03 |
| DQ164203 | TD02 / TD03 |
| DQ164204 | TD02 / TD03 |
| DQ164205 | TD02 / TD03 |
| DQ164206 | TD02 / TD03 |
| DQ176637 | TD02 / TD03 |
| DQ211652 | TD02 / TD03 |
| DQ256376 | TD02 / TD03 |
| DQ374650 | TD02 / TD03 |
| DQ374651 | TD02 / TD03 |
| DQ374652 | TD02 / TD03 |
| DQ374653 | TD02 / TD03 |
| DQ377178 | TD02 / TD03 |
| DQ377179 | TD02 / TD03 |
| DQ377180 | TD02 / TD03 |
| DQ411029 | TD02 / TD03 |
| DQ411030 | TD02 / TD03 |
| DQ411031 | TD02 / TD03 |
| DQ411032 | TD02 / TD03 |
| DQ411033 | TD02 / TD03 |
| DQ411034 | TD02 / TD03 |
| DQ411035 | TD02 / TD03 |
| DQ431693 | TD02 / TD03 |
| DQ431694 | TD02 / TD03 |
| DQ431695 | TD02 / TD03 |
| DQ431696 | TD02 / TD03 |
| DQ431697 | TD02 / TD03 |
| DQ431698 | TD02 / TD03 |
| DQ431699 | TD02 / TD03 |
| DQ431700 | TD02 / TD03 |
| DQ431701 | TD02 / TD03 |
| DQ431702 | TD02 / TD03 |
| DQ431703 | TD02 / TD03 |
| DQ431704 | TD02 / TD03 |
| DQ431705 | TD02 / TD03 |
| DQ431706 | TD02 / TD03 |
| DQ431707 | TD02 / TD03 |
| DQ431708 | TD02 / TD03 |
| DQ431709 | TD02 / TD03 |
| DQ431710 | TD02 / TD03 |
| DQ431711 | TD02 / TD03 |
| DQ431712 | TD02 / TD03 |
| DQ666448 | TD02 / TD03 |
| DQ666449 | TD02 / TD03 |
| DQ666450 | TD02 / TD03 |
| DQ666451 | TD02 / TD03 |
| DQ666452 | TD02 / TD03 |
| DQ786572 | TD02 / TD03 |
| DQ786573 | TD02 / TD03 |
| EU249803 | TD02 / TD03 |
| FJ483548 | TD02 / TD03 |
| FJ483549 | TD02 / TD03 |
| FJ527738 | TD02 / TD03 |
| FJ766331 | TD02 / TD03 |
| FJ766332 | TD02 / TD03 |
| GQ379157 | TD02 / TD03 |
| GQ379158 | TD02 / TD03 |
| GQ379159 | TD02 / TD03 |
| GQ379160 | TD02 / TD03 |
| GQ379161 | TD02 / TD03 |
| GQ851602 | TD02 / TD03 |
| GQ851603 | TD02 / TD03 |
| GQ851604 | TD02 / TD03 |
| GQ851605 | TD02 / TD03 |
| GQ851606 | TD02 / TD03 |
| GQ851607 | TD02 / TD03 |
| GQ851608 | TD02 / TD03 |
| GU011992 | TD02 / TD03 |
| GU827998 | TD02 / TD03 |
| GU827999 | TD02 / TD03 |
| GU828000 | TD02 / TD03 |
| GU828001 | TD02 / TD03 |
| GU828002 | TD02 / TD03 |
| GU828003 | TD02 / TD03 |
| GU828004 | TD02 / TD03 |

**Table S4.**
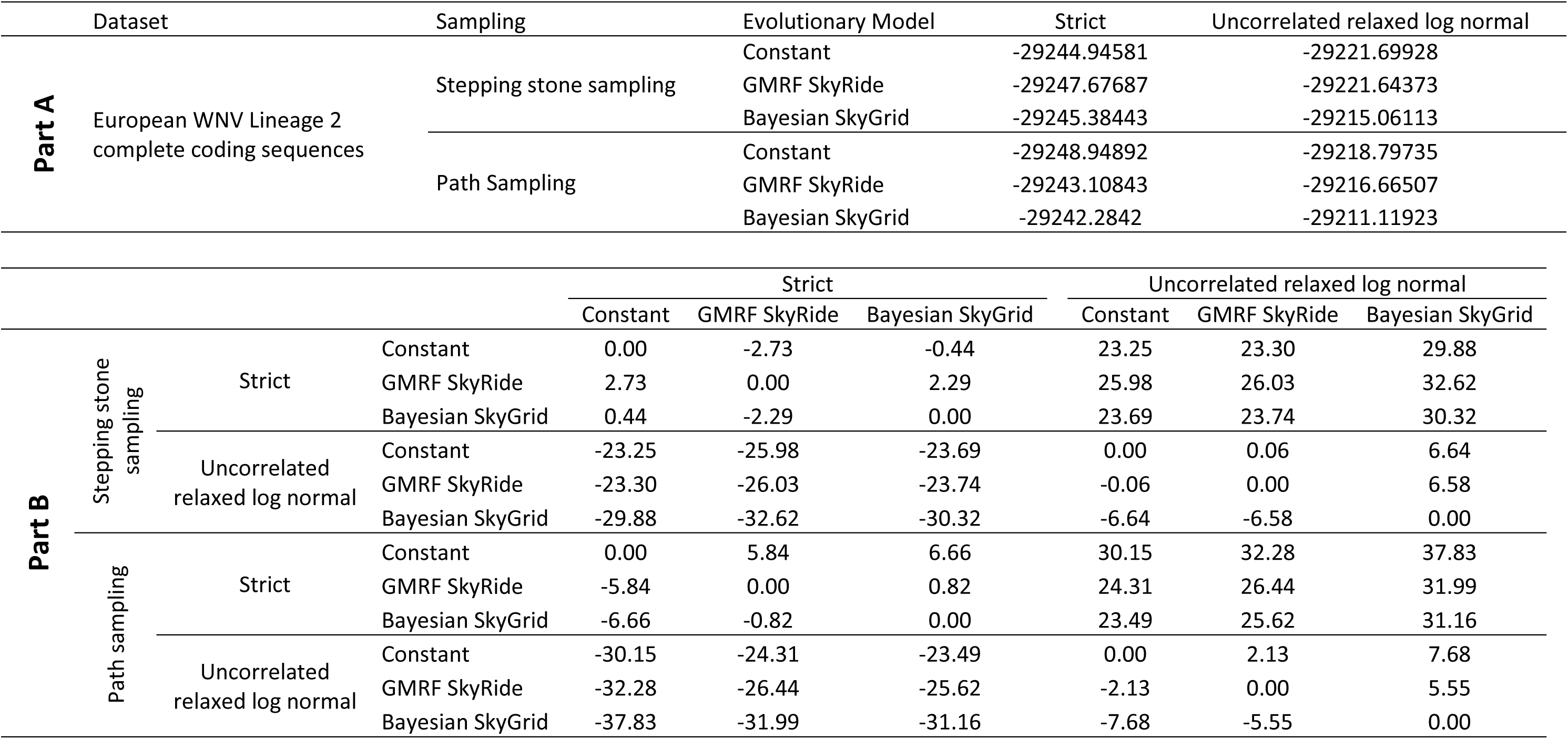
Summary and comparison of parameter values from Beast analysis, parts A and B (A) Result of marginal likelihood (log) estimation path sampling and stepping stone sampling methods for West Nile Virus Lineage 2 dataset using different coalescent models, and strict and uncorrelated relaxed log normal molecular clock models. (B) Calculation of best coalescent model and molecular clock model using Bayes factor. Bayes factor range 1-3 means hardly worth mentioning, 3- 20 means postive support, 20-150 means strong support and >150 overwhelming support.

